# Impairments in the early consolidation of spatial memories via group II mGluR agonism in the mammillary bodies

**DOI:** 10.1101/2023.12.13.571542

**Authors:** Michal M. Milczarek, James C. Perry, Eman Amin, Salma Haniffa, Thomas Hathaway, Seralynne D. Vann

**Affiliations:** School of Psychology & Neuroscience and Mental Health Innovation Institute, Cardiff University, Cardiff, CF10 3AT

## Abstract

mGluR2 receptors are widely expressed in limbic brain regions associated with memory, including the hippocampal formation, retrosplenial and frontal cortices, as well as subcortical regions including the mammillary bodies. mGluR2/3 agonists have been proposed as potential therapeutics for neurological and psychiatric disorders, however, there is still little known about the role of these receptors in cognitive processes, including memory consolidation. To address this, we assessed the effect of the mGluR2/3 agonist, eglumetad, on spatial memory consolidation in both mice and rats. Using the novel place preference paradigm, we found that post-sample injections of eglumetad impaired subsequent spatial discrimination when tested 6 hours later. Using the immediate early gene c-*fos* as a marker of neural activity, we showed that eglumetad injections reduced activity in a network of limbic brain regions including the hippocampus and mammillary bodies. To determine whether the systemic effects could be replicated with more targeted manipulations, we performed post-sample infusions of the mGluR2/3 agonist 2*R*,4*R*-APDC into the mammillary bodies. This impaired novelty discrimination on a place preference task and an object-in-place task, again highlighting the role of mGluR2/3 transmission in memory consolidation and demonstrating the crucial involvement of the mammillary bodies in post-encoding processing of spatial information.

## Introduction

Memory formation is a complex process, involving encoding, consolidation, and recall. These processes depend on a network of brain regions whose relative contribution varies according to memory stage as well as its content. The medial diencephalon, including the mammillary bodies, has long been implicated in memory processing^[1–3]^, especially in relation to encoding^[4–7]^. By contrast, less is known about its possible contributions to consolidation, which has been more closely aligned with the medial temporal lobe and cortical regions^[8,9]^, which are sites directly involved in memory storage. However, increasing evidence suggests an active role for subcortical regions in orchestrating systems-wide memory consolidation, via neuromodulatory inputs^[10,11]^ and generation of oscillatory activity^[12]^. This role may be extended to the mammillary bodies given their importance for arousal and coordinating hippocampal-network activity^[2,13]^.

The mammillary bodies form part of an extended memory system (or the circuit of Papez), receiving their major glutamatergic input from the subiculum via the postcommissural fornix and projecting to the anterior thalamic nuclei, which in turn reciprocally connect with the hippocampal formation and cortical regions involved in mnemonic processing^[14]^. Mammillary body activity is modulated via their reciprocal connections with Gudden’s tegmental nuclei, and inputs from the septum and supramammillary nuclei^[2]^. This enables the mammillary bodies to modulate and amplify relevant signals for subsequent processing in downstream regions^[2,13]^, in addition, their role in the propagation of sharp-wave ripple-related activity^[15]^ provides a further mechanism via which the mammillary bodies may contribute to consolidatory processes^[16]^.

The mammillary bodies show evidence of highly specialized glutamatergic transmission: they express vesicular glutamate transporters as well as ionotropic and metabotropic glutamate receptors^[17,18]^. Indeed, the medial mammillary nucleus has some of the highest metabotropic glutamate receptor 2 (mGluR2) expression levels, along with other regions in the Papez circuit including the hippocampus, and retrosplenial cortex^[19]^. The distribution of mGluR2 in the brain might suggest that modulating mGluR2 activity would impact memory processes. However, the effects of mGluR2/3 agonists on spatial working memory tasks appear mixed^[20]^, with some studies finding impairments^[21,22]^ while others showing improvements^[23]^ and others showing no effect on cognition^[24]^. However, metabotropic glutamate receptors in general have also been implicated in memory consolidation^[25]^, due to their role in slower, longer-lasting modulatory effects^[26]^, including long-term depression^[27]^, and mGluR2 could therefore be mediating consolidatory processes within the Papez circuit^[28]^.

To assess the effects of mGlu2 receptor manipulation on early consolidation processes, rats and mice were systemically injected with an mGluR2/3 agonist, eglumetad (LY354740), immediately after the sample phase of a novel place preference task and a discrimination test was carried out after a 6-hour delay. A motility test was used to determine whether the injections had any overall effects on activity. To identify which brain regions were affected by systemic eglumetad injections, we quantified the expression of the immediate early gene c-*fos* across the Papez circuit of mice that had explored a novel environment after receiving either an injection of eglumetad or saline.

The next step was to determine whether the systemic effects of mGluR2/3 agonists could be replicated with more targeted administration and for this we looked at the effects of mGlu2 receptor agonism in the mammillary bodies. The mammillary bodies were selected as they have particularly high levels of mGlu2 receptor expression^[19]^ and slice studies have shown the mGluR2/3 agonist, APDC, to inhibit mammillary body activity^[29]^. Animals were tested on two spatial discrimination tasks that assess incidental learning and rely on an animal’s innate preference for novelty^[3]^. By looking at spatial discrimination behavior with a 6-hour delay, and administering the drug post-sample, it was possible to study the impact of an mGluR agonist on early consolidation while leaving the animals unaffected by any acute effects of the drug during both encoding and recall periods.

## Results

### Experiment 1

We first tested the impact of eglumetad injections in rats on open field exploration and found no effect on total distance travelled or mean distance from the center of the arena at 30 min post-injection (total distance travelled: t = 1.41, *p =* 0.090, d = 0.37; mean distance from center: t = -0.98, *p =* 0.84, d = -0.26; **Fig. 1 a-c)** or at 6 h post-injection (total distance travelled: t = -0.34, *p =* 0.63, d = -0.09; mean distance from center: t = 0.09, *p =* 0.46, d = 0.09, **Fig. S1 a-c**; one-sided within-subject permutation tests).

**Figure 1.**
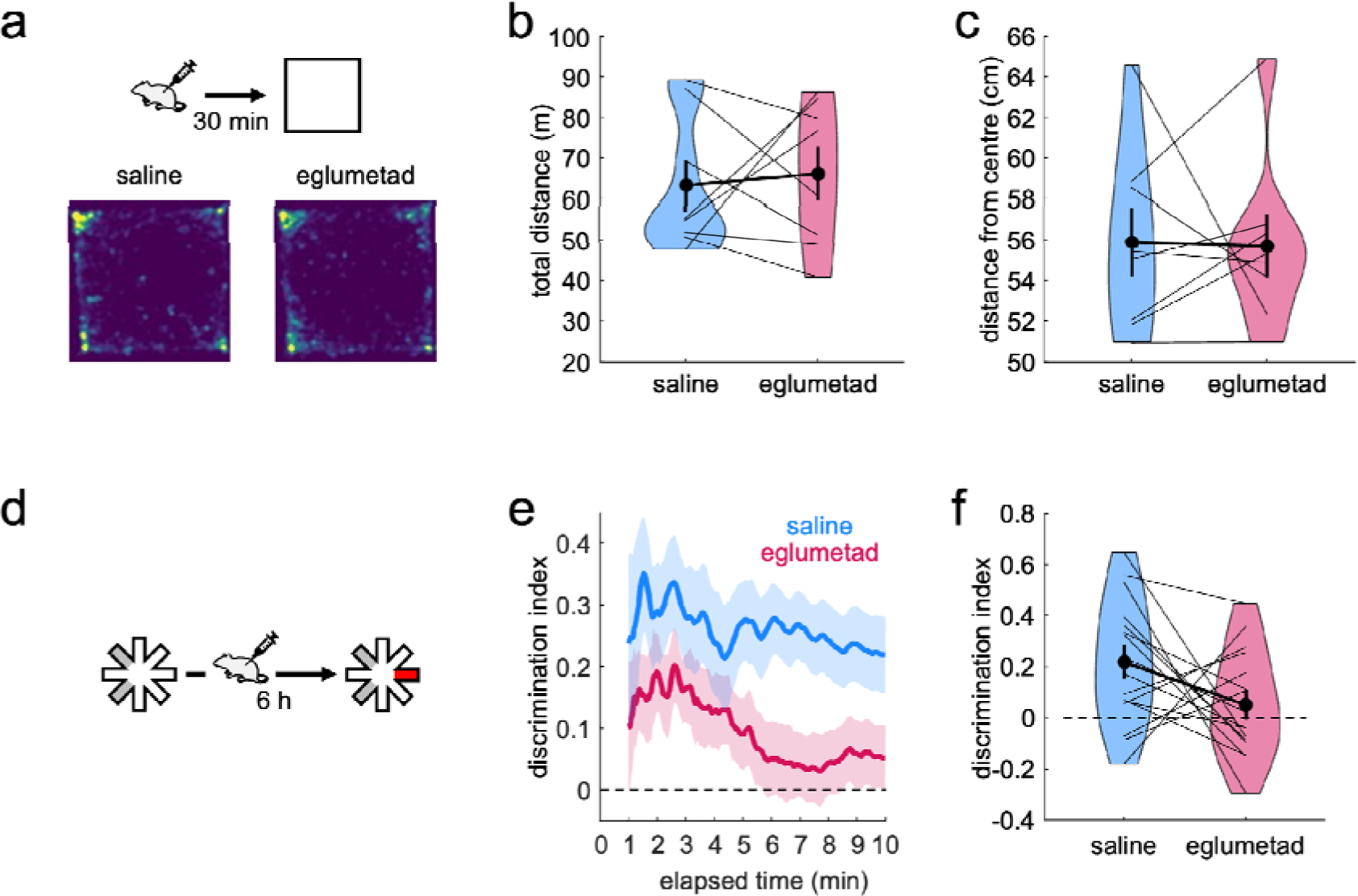
The effect of eglumetad on open field activity and novel place preference in rats. **a** – Schematic representation of the open field task and heatmaps representing mean arena occupancy (n = 8, within-subject design). **b** – Violin plot of total distance travelled. The vertical bars are standard error of the mean, black dots represent the means, thin lines connect data from individual animals and the boldened line connects the means. **c** – Violin plot of the mean distance from center of the open field. Lower values suggest less anxiety. **d** – Schematic representation of the novel place preference task (n = 16, within-subject design): rats were presented with two sample maze arms for 10 min, followed by intraperitoneal injections and 6 h later given access to three arms (1 novel and 2 familiar) for 10 min. The discrimination index at test was calculated as the ratio of the difference in time spent in the novel and familiar arms and the summed time spent in the novel and familiar arms. **e** – Timeline of the cumulative discrimination index following saline or eglumetad injections. The solid lines represent mean values and the shaded areas, the standard error of the mean. **f** – Violin plot of mean discrimination indices, calculated over 10 min of maze exploration at test. The horizontal line in **e** and **f** denotes chance-level performance.

Next, we investigated the effect of the drug on memory consolidation using the novel place preference (NPP) task with a 6 h delay between sample and test. Post-sample injections of eglumetad impaired novel spatial discrimination (**Fig. 1 d**), as demonstrated by a significant reduction in cumulative discrimination scores when compared with saline injections (t = 1.79, *p =* 0.047, d = 0.55; one-sided within-subject permutation test; **Fig. 1 e&f**). Furthermore, rats displayed clear novel place preference with above-chance cumulative discrimination scores following saline injections (t = 3.59, *p =* 1.62×10^-3^; one-sided within-subject permutation test) whereas they performed at chance following eglumetad injections (t = 1.02, *p =* 0.16). The differences in discrimination were not driven by changes in general activity as there was no effect of drug on total number of arm entries or mean duration of time spent in the arms either at sample or test phases of the task (number of entries at sample: t = -0.97, *p =* 0.38, d = -0.21; mean arm visit time at sample: t = 0.63, *p =* 0.54, d = -0.14; number of entries at test: t = 0.06, *p =* 1, d = 0.012; mean arm visit time at sample: t = 0.68, *p =* 0.51, d = -0.15; two-sided within-subject permutation tests). However, we found that total active exploration at sample was negatively correlated with subsequent discrimination scores at test following saline injections (r = -0.56, *p =* 0.024) but not following eglumetad injections (r = -0.04, *p =* 0.88). We therefore tested if inclusion of sample total exploration data improved the estimation of discrimination scores at test and found that it marginally did (λ = 3.88, *p =* 0.049, compared to a model lacking sample data), while still returning the main effect of drug (F_1,29_ = 5.45, *p =* 0.027) and a trend for sample total exploration on D2 scores at test (F_1,29_ = 4.13, *p =* 0.051).

We also measured the concentration of the drug in rat brains at increasing intervals between 10 and 360 min from injection, confirming its presence immediately after injection and a near full wash-out by 6 h (5.5-fold decrease, **Fig. S2**).

### Experiment 2

As with rats (**Experiment 1**), eglumetad injections had no effect on the total distance travelled in the open field in mice (t = 0.44, *p =* 0.33, d = 0.44; one-sided between-subject permutation test; **Fig. 2 a&b**). However, unlike the effect on rats, the injections in mice resulted in greater exploration in the center of the arena, revealing an anxiolytic effect of the drug (t = 2.59, *p =* 9.93×10^-3^, d = 1.03, one-sided between-subject permutation test; **Fig. 2 a&c**), likely driven by the higher dose used in mice than rats (10mg/kg vs 6mg/kg, respectively).

**Figure 2.**
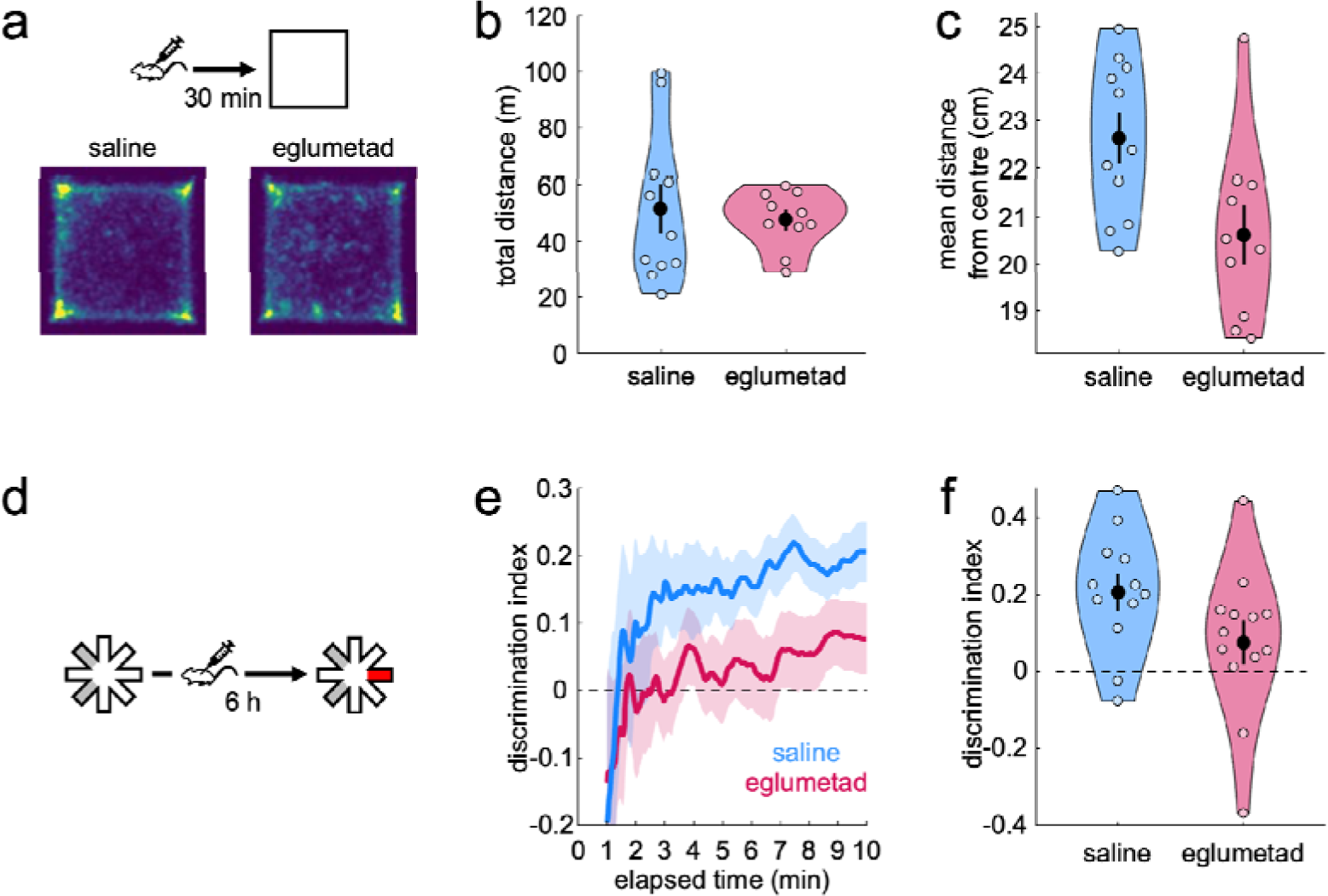
The effect of eglumetad on open field activity and novel place preference in mice. **a** – Schematic representation of the open field task and heatmaps representing mean arena occupancy (saline: 7 female, 4 male; eglumetad: 4 female, 6 male). **b** – Violin plot of total distance travelled, as in **Fig. 1 b**. **c** – Violin plot of the mean distance from center of the open field. Lower values signify less anxiety. **d** – Schematic representation of the novel place preference task. **e** – Timeline of the cumulative discrimination index in mice that received saline (5 female, 7 male, blue) or eglumetad (5 female, 8 male, red) following sample. The solid lines represent mean values and the shaded areas, the standard error of the mean. **f** – Violin plot of mean discrimination indices, calculated over 10 min of maze exploration at test. The horizontal line in **e** and **f** denotes chance-level performance.

Eglumetad affected performance on the NPP task in mice (**Fig. 2 d**) in the same way as in rats, with saline-injected mice displaying above-chance discrimination (t = 4.6, *p =* 9.77×10^-4^) and eglumetad-injected mice showing no preference (t = 1.43, *p =* 0.090; one-sided between-subject permutation tests, **Fig. 2 e-f**). This was reflected in significant drug-related differences in discrimination scores (t = 1.85, *p =* 0.037, d = 0.69, one-sided between-subject permutation test). The impaired discrimination following eglumetad injections did not reflect changes in overall activity as there was no effect of drug on the number of arm entries and the mean duration of time spent in the arms at either sample or test phases of the task (number of entries at sample: t = -1.19, *p =* 0.23, d = -0.44; mean arm visit time at sample: t = 2.07, *p =* 0.050, d = -0.77; number of entries at test: t = 0.016, *p =* 0.98, d = 0.006, mean arm visit time at test: t = -0.45, *p =* 0.69, d = 0.17, two-sided permutation tests). As with rats, we found total sample exploration time to negatively correlate with subsequent discrimination at test, this time both for saline (r = -0.58, *p =* 0.0490) and eglumetad (r = -0.56, *p =* 0.0495). Inclusion of sample exploration data significantly improved the mixed effects model (λ = 7.91, *p =* 0.0049), still returning a main effect of drug (F_1,22_ = 9.33, *p =* 0.0058) and a main effect of total exploration (F_1,22_ = 9.3, *p =* 0.0059). Factoring in sex did not improve the model (λ = 0.007, *p =* 0.92), suggesting the effect of the drug was equivalent in males and females.

### Experiment 3

To identify which brain areas were affected by systemic eglumetad injections (10 mg/kg), we looked at the effects of the drug on novelty-induced *c-fos* expression across limbic brain regions in mice. Mice were injected with either eglumetad or saline 30 mins before being placed in a novel environment. While eglumetad injections had no effect on motility (t = 0, *p =* 0.68, d = -0.23; one-sided between-subject permutation test; **Fig. 3 a&b**), *c-fos* levels showed a marked reduction across multiple limbic areas. A mixed effect model (‘Density ∼ Drug × Region +(1|Animal)’) returned a main effect of drug (F_1,11_ = 9.32, *p =* 2.72×10^-3^), a main effect of region (F_1,11_ = 13.94, *p =* 4.86×10^-20^) and a drug × region interaction (F_14,154_ = 1.83, *p =* 0.039; **Fig. 3 c&d**). Pairwise comparisons revealed significant differences in all regions of interest apart from the visual cortex, postsubiculum, anteroventral and anteromedial thalamic nuclei (**Fig. S3**).

**Figure 3.**
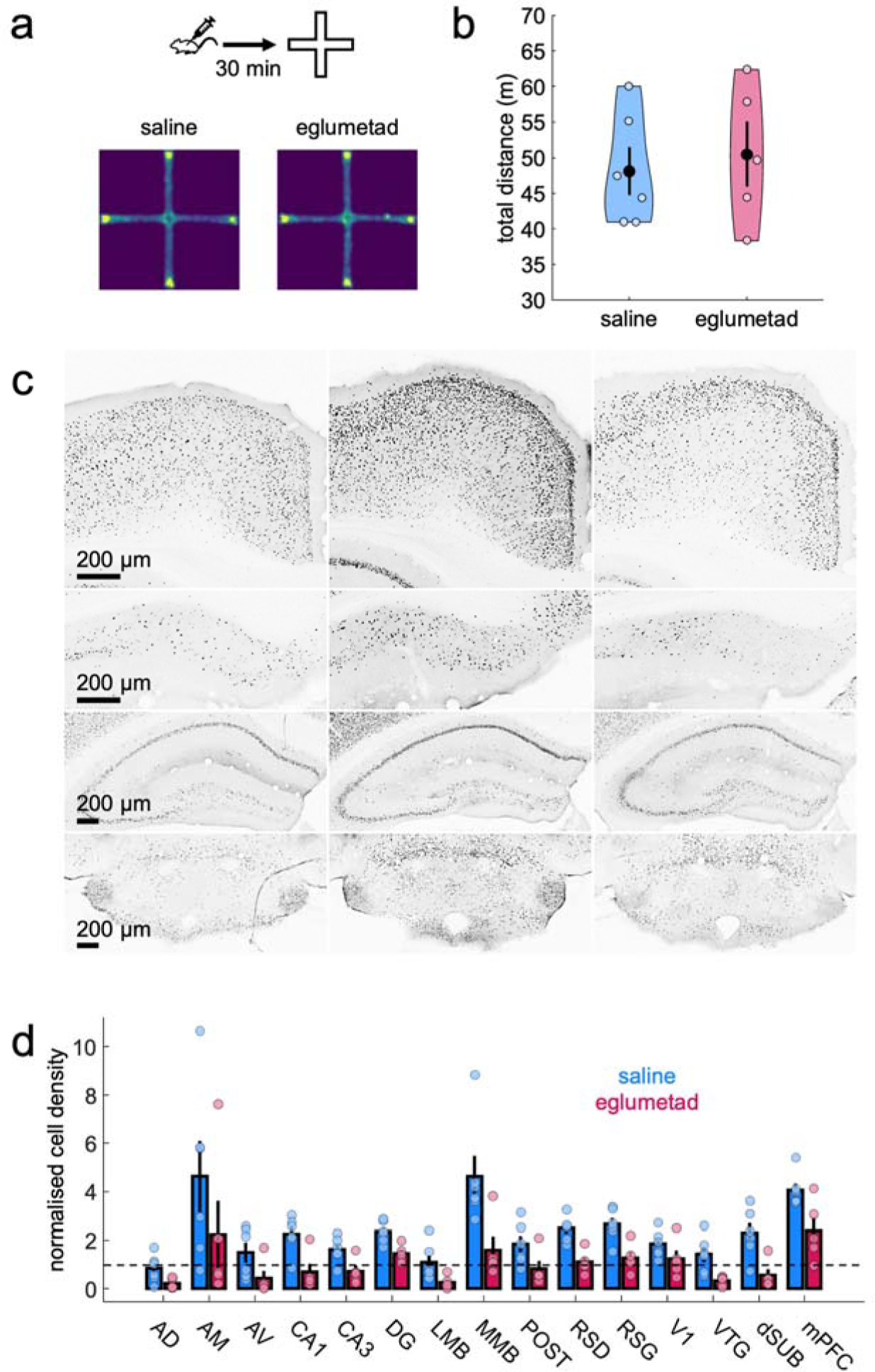
The effect of eglumetad on motility and *c-fos* induction in mice (Experiment 3). **a** – Heatmaps showing cross-maze motility following saline or eglumetad (10 mg/kg) injections. **b** – Violin plots representing total distance travelled in cross-maze. Large black dots are mean values, the error bars are standard error of the mean and smaller light dots are data from individual animals (saline: n = 6, eglumetad: n = 5). **c** – Representative images of selected areas of the Papez circuit (from top to bottom: retrosplenial cortex, dorsal subiculum, dorsal hippocampus, mammillary bodies) displaying differences in *c-fos* staining in home-cage controls (first column), mice under saline (second column) and mice under eglumetad (third column). Images from other regions as well as the quantification of home-cage control *c-fos* densities are displayed in **Fig. S3**. **d** – Bar chart showing mean regional differences in relative *c-fos* expression (fold density change relative to mean home-cage control values) with counts from mice under saline or eglumetad in blue and red, respectively. The error bars are standard error of the mean and data from individual animals are shown as colored dots. The horizontal line represents no change relative to home-cage control. Abbreviations: AD – anterodorsal thalamic nucleus, AM – anteromedial thalamic nucleus, AV – anteroventral thalamic nucleus, CA1/CA3 – hippocampal subfields, DG – dentate gyrus, LMB – lateral mammillary bodies, MMB – medial mammillary bodies, POST – postsubiculum, RSD – dysgranular retrosplenial cortex, RSG – granular retrosplenial cortex, V1 – primary visual cortex, VTG – ventral tegmental nucleus of Gudden, dSUB – dorsal subiculum, mPFC – medial prefrontal cortex.

### Experiment 4

Experiment 3 identified several limbic regions that were affected by eglumetad injections. Among these, the mammillary bodies show high levels of circumscribed mGluR2 expression (**Fig. 4 e**) and they have also been shown to directly respond to group II metabotropic agonism^[29]^. As such, we hypothesized that local infusion of mGluR2/3 agonists into the mammillary bodies would be sufficient to impair spatial memory consolidation.

**Figure 4.**
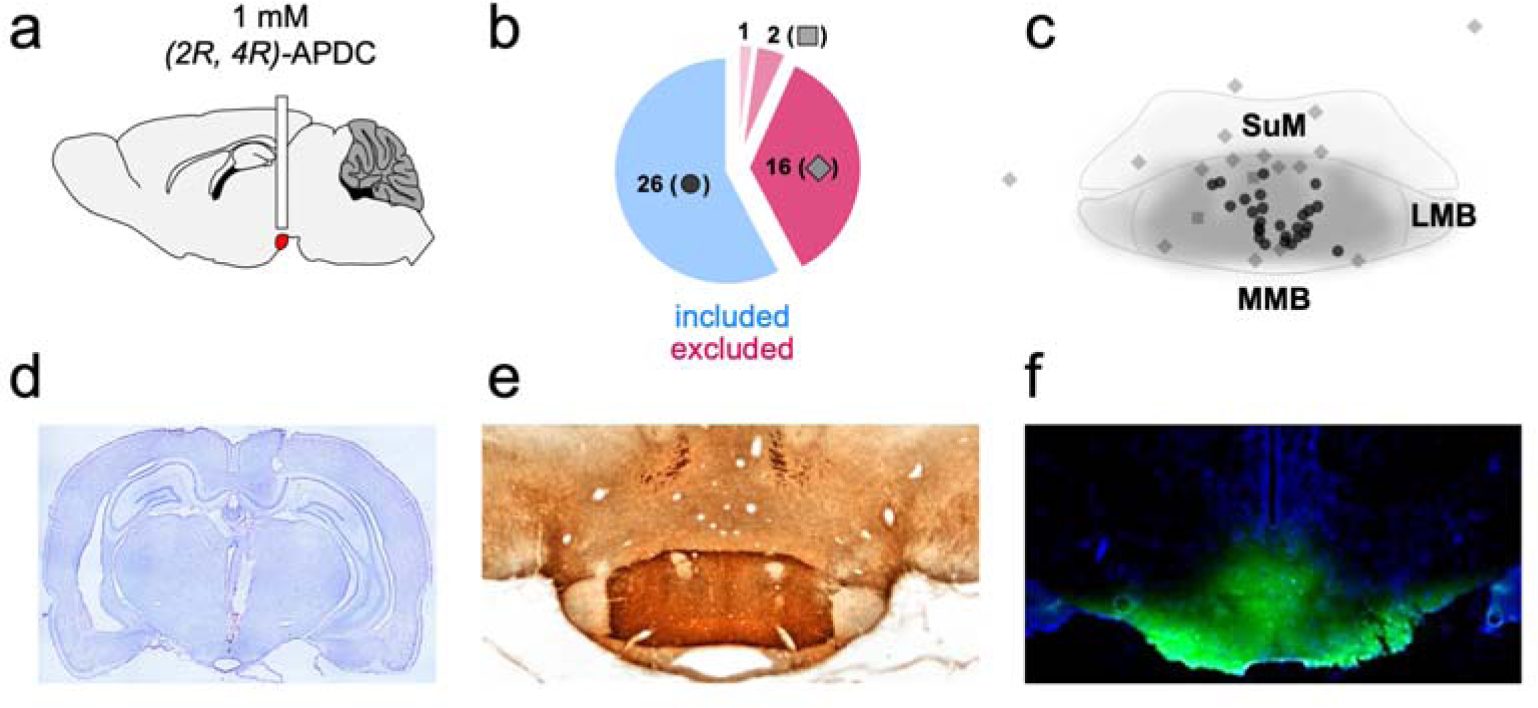
Local drug infusion in the mammillary bodies. **a** – Diagrammatical representation of the guide cannula implant. The mammillary bodies are highlighted in red. **b** – Subject classification based on histological verification. Out of 45 cannulation surgeries, 26 were classified as successful and 16, unsuccessful. Two animals with good cannula placement were rejected due to excess damage and one animal was removed from the study due to unrelated health issues. **c** – Estimated cannula tip positions projected onto a mean mammillary body outline. Note that the average boundaries of the mammillary bodies are only approximated here, see **Fig. S4** for individual coronal levels as well as images of cannulated mammillary bodies for all cases. Dots represent successful canulations, diamonds – misplaced cannulations and squares, excess damage (in both **b&c**). SuM – supramammillary nucleus, LMB – lateral mammillary body, MMB – medial mammillary body. **d** – Coronal section (Nissl staining) displaying the guide cannula track and infusion cannula tip location in the mammillary bodies. **e** – mGluR2 immunostaining in the mammillary bodies. The label clearly differentiates the medial mammillary nucleus from the surrounding tissue, including the supramammillary nucleus. **f** – the spread of locally infused fluorescent dye (acridine orange) in the mammillary bodies.

To test this, we cannulated 45 rats to locally infuse the mGluR2/3 agonist, *2R*, *4R*-APDC (**Fig. 4 a**). Postmortem validation, including cresyl (**Fig. 4 d**) and DAB staining (**Fig. S4 b** for individual subjects), and in some cases, infusions of a fluorescent dye (**Fig. 4 f**), confirmed correct cannula placement, with limited tissue damage, in 26 rats (**Fig. 4 b-c** & **Fig. S4 a**). To ensure local infusions of APDC had no effect on motility, the rats were tested on open field exploration following saline and APDC infusions. There was no effect of drug on total distance travelled (t = -1.01, *p =* 0.83, d = -0.26) or mean distance from the center of the arena (t = 1.5, *p =* 0.090, d = 0.39, one-sided within-subject permutation tests; **Fig. 5 a-c**).

**Figure 5.**
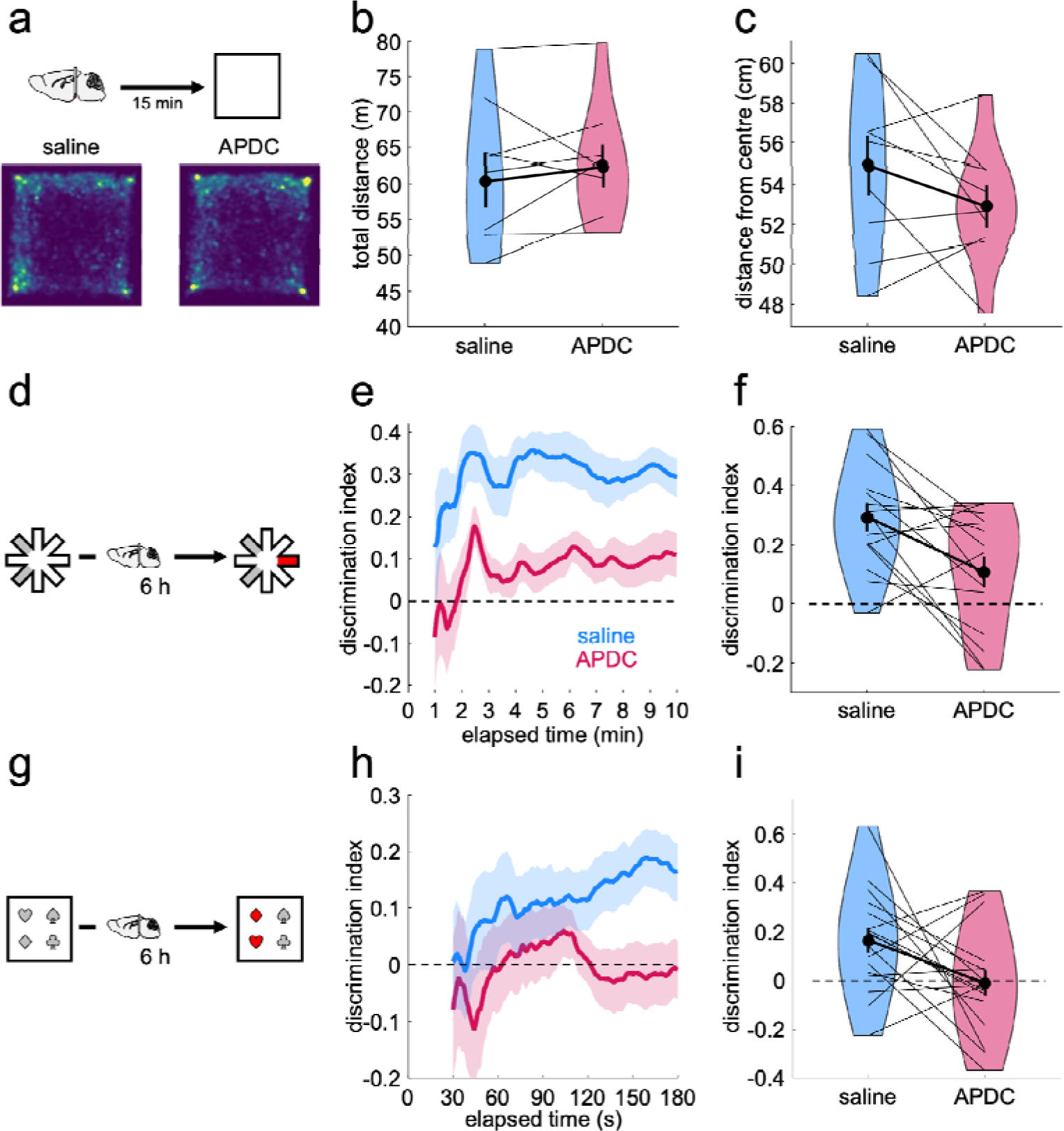
The effect of APDC infusions into the mammillary bodies on open field activity, novel place preference and object-in-place in rats. **a** – Schematic representation of the open field task and heatmaps representing mean arena occupancy (within-subject n=9). **b** – Violin plot of total distance travelled, as in Fig. 1 b. **c** – Violin plot of the mean distance from center of the open field. Lower values suggest less anxiety. **d** – Schematic representation of the NPP task. **e** – Timeline of the cumulative discrimination index in rats that received saline (blue) or APDC infusions (red) following sample (within-subject design, n=15). The solid lines represent mean values and the shaded areas, the standard error of the mean. **f** – Violin plot of mean discrimination indices, calculated over 10 min of maze exploration at test. **g** – Schematic representation of the object-in-place task. **h** – Timeline of the cumulative discrimination index in rats that received saline (blue) or APDC infusions (red) following sample (within-subject design, n=17). The solid lines represent mean values and the shaded areas, the standard error of the mean. **i** – Violin plot of mean discrimination indices, calculated over 10 min of maze exploration at test. The horizontal line in e, f, g, i denotes chance-level performance.

For the NPP task, we found a similar pattern to that of the systemic injection experiments, with APDC infusions leading to reduced discrimination (t = 3.06, *p =* 5.08×10^-3^, d = 0.54, one-sided within-subject permutation test). However, unlike the systemic injections, rats’ performance was above chance following both saline (t = 6.38, *p =* 6.1×10^-5^) and APDC infusions (t = 2.07, *p =* 0.03; one-sided within-subject permutation tests), showing rats could discriminate the novel arm under both drug conditions. The main effect of drug was not driven by differences in overall activity levels at either sample or test (number of entries at sample: t = -0.32, *p =* 0.81, d = -0.074; mean arm visit time at sample: t = 1.08, *p =* 0.30, d = -0.24; number of entries at test: t = -0.20, *p =* 0.89, d = -0.05, mean arm visit time at test: t = -0.12, *p =* 0.91, d = 0.028; two-sided within-subject permutation tests). There were no relationships between sample exploration times and discrimination at test (saline: r = 0.0033, *p =* 0.99, APDC: r = 0.09, *p =* 0.74) and inclusion of sample data did not improve a linear mixed effect model (λ = 0.22, *p =* 0.64).

To test the specificity of these effects, we looked at performance in the rats excluded due to incorrect cannula placements. Rats were able to discriminate the novel location both after saline (t = 2.49, *p =* 0.017) and APDC infusions (t = 2.59, *p =* 0.016) and there was no difference in discrimination scores across drug conditions (t = -0.43, *p =* 0.67, d = -0.15; **Fig. S5**).

Finally, we tested whether the effect of local infusions on the consolidation of spatial memory could be generalized to another novelty-driven task, the object-in-place task (**Fig. 5 g**). We found that rats were able to discriminate the displaced objects at test following saline infusions (t = 3.27, *p =* 2.22×10^-3^, one-sided within-subject permutation test), whereas rats performed at chance following APDC infusions (t = -0.18, *p =* 0.57, one-sided within-subject permutation test). These drug-related differences were reflected in a significant difference in discrimination scores (t = 2.13, *p =* 0.024, d = 0.46, one-sided within-subject permutation test). As in the NPP task, this difference was not due to changes in general exploratory behaviors at sample or at test (sample: t = 0.79, *p =* 0.44, d = 0.17; test: t = 1.31, *p =* 0.21, d = 0.28; two-sided within-subject permutation tests). While total sample exploration did not relate to discrimination scores at test (saline: r = -0.06, *p* = 0.81; APDC: r = -0.10, *p* = 0.70; model with sample exploration versus simpler model: λ = 0.22, *p =* 0.64), object preference at sample (sample discrimination scores for subsequently displaced objects) showed a positive relationship with test discrimination scores for rats infused with saline (r = 0.67, *p =* 0.0035) but not with APDC (r = -0.33, *p =* 0.20). Consistent with this, an interaction model (sample discrimination scores x drug) offered a better fit compared to a simple model (λ = 10.73, *p =* 0.0047) and to an additive model (λ = 9.77*, p =* 0.0018), returning a significant drug x sample_D2 interaction (F_1,30_ = 11.31, *p =* 0.0021) and a main effect of drug (F_1,30_ = 7.71, *p =* 0.0094).

## Discussion

We identified a role for the mGluR2/3 agonist, eglumetad, in early spatial memory consolidation and this was mediated, in part, by its effect on the medial mammillary bodies. Post-sample administration of mGlu2 receptor agonists either injected systemically or infused into the mammillary bodies impaired performance on a novel place preference task. The systemic effects were found to be highly reliable as the same pattern was observed across both mice and rats; in mice the effects were observed in both males and females. mGlu2 receptors are strongly expressed in the Papez circuit in the rodent brain^[19]^, a brain network important for memory^[2]^. Consistent with this, we found that systemic injections of eglumetad in mice markedly reduced novelty-induced c-*fos* expression across the Papez circuit, including the mammillary bodies, hippocampus, retrosplenial cortex and ventral tegmental nucleus of Gudden, indicating an inhibitory effect across this memory network. This pattern of reduced activity replicated findings from a previous glucose imaging experiment^[30]^, establishing the impact of mGluR2/3 agonists on limbic memory networks. To determine whether local inhibition of the mammillary bodies could produce a similar behavioral effect to a systemic mGluR2/3 agonist, or whether network-wide changes were necessary, we selectively targeted the mammillary bodies using the mGluR2/3 agonist, 2*R*, 4*R*-APDC. A remarkably consistent pattern of findings was observed with the local manipulation as with the systemic injections, suggesting the systemic effect of mGluR2/3 agonism was in part driven by its effect on the mammillary bodies. Furthermore, we extended our findings by showing impaired consolidation of a separate behavioral task, the object-in-place task, which is known to be sensitive to lesions of mammillary body pathways^[3]^.

Neither systemic administration of an mGluR2/3 agonist, nor selective infusion into the mammillary bodies, affected the overall activity of rats or mice, as measured by open field exploration. Mice did, however, explore the center of the arena more following eglumetad injections, consistent with the anxiolytic effect of mGluR2/3 agonists^[31]^. Mammillary body lesions have also been found to reduce anxiety on the open field task ^[31]^, although we did not find increased center occupancy in locally infused rats, or in systemically injected rats, which may reflect the amount of drug administered. There was also no difference in animals’ overall activity in the test phase of the discrimination tasks, despite impaired performance. This would suggest that the discrimination effects are driven by memory impairments resulting from impoverished consolidation, rather than non-specific changes to activity during the test stage. Consistent with this, the brain occupancy data showed that following systemic administration, eglumetad had mostly cleared from the brain after the first hour. As such, the post-sample drug manipulations can be considered in terms of their impact on early consolidation rather than directly affecting the recall stage.

We observed no mean differences in sample activity, as would be expected given the manipulations were carried out after the sample phase. However, we did find relationships between sample activity and test scores at an individual animal level across most of the behavioral tasks, likely driven by animals’ biases towards certain objects or maze arms. By including these sample exploration data in the analyses, we were able to reduce some of the variance and better isolate the effect of experimental treatment, making the data more robust and highlighting the potential benefit of using sample data in these types of experiments. The current study employed a two-phase design with a 6-hour consolidation window, which capitalized on rats’ and mice’s natural ultradian activity patterns. It is unclear whether manipulating the length of the sample-test delay would have produced an equivalent deficit, however, lesions of the mammillary bodies or their principal efferent pathway have been shown to impair performance on the OIP task at shorter delays ^[3,32]^. Considering the systems-wide impact of eglumetad injections on *c-fos* expression, it might be predicted that that memory storage over longer delays, requiring greater cortical engagement, would also be vulnerable to mGluR2/3 agonist effects.

Almost all previous behavioral studies assessing the role of the mammillary bodies have involved permanent lesions, making it difficult to draw conclusions about their contributions to specific stages of memory processing. One of the few inactivation studies found that manipulation of the mammillary bodies in rabbits immediately after trace conditioning impaired reflex learning. By contrast, there was no effect if inactivation occurred 3 hours after training, suggesting a role for the mammillary bodies during the early consolidation period^[33]^. Glucose imaging and cytochrome oxidase imaging studies have also shown increased mammillary body activity during the first few hours following task learning^[34–36]^, consistent with a role for the mammillary bodies in this early post-encoding period.

Temporary inactivation of both dorsal subiculum^[37]^ and the anterior thalamic nuclei^[38]^, regions that send inputs to, and receive outputs from the mammillary bodies, respectively, have both been shown to affect memory consolidation, suggesting that they, along with the mammillary bodies, form part of an extended consolidation network. Indeed, the mammillary bodies’ role in mediating sharp-wave ripple-related activity between the dorsal subiculum and anterior thalamic nuclei might be key to this role in consolidation^[15,39]^. However, there are also high expression levels of mGluR2 in the ventral tegmental nucleus of Gudden and, in agreement with this, we found reduced c-*fos* expression in this region following eglumetad injections. Therefore, the reciprocal mammillary body-tegmental nuclei pathway could also be contributing to consolidatory mechanisms, and the behavioral impairments observed.

mGluR2/3 agonists have been identified as a potential treatment for schizophrenia^[40]^. The present results are difficult to interpret in terms of therapeutics since we investigated only the acute effects of drug administration. Chronic administration may produce different results on memory consolidation, with the agonist present at multiple stages of the task. Furthermore, downregulation of receptors and compensatory responses following chronic administration could also produce different effects. Nevertheless, future work into mGluR2-targeting agents as a treatment should consider the possible impact on longer-term memory processes.

Improving the therapeutic potential of mGluRs requires mechanistic understanding of their roles across multiple brain systems. Our results demonstrate their relevance to long-term memory via interactions within the limbic system and highlight the specific role of the mammillary bodies in early memory consolidation. The effect of systemic eglumetad delivery on *c-fos* expression suggests that the effects on consolidation may be driven by overall depression of neuronal activity. In support of this, group II mGluRs act as glutamate autoreceptors whose activation results in reduced presynaptic glutamate release, as seen in the hippocampus, prefrontal cortex, sensory cortices and others^[41,42]^. In the case of the mammillary bodies, mGluR2 is, however, likely to directly exert its function by reducing neuronal excitability postsynaptically^[29]^ (although the contribution of presynaptic and astrocytic mGluR2 cannot be ruled out). As such, impaired early consolidation following local drug infusion may simply reflect diminished mammillary body activity levels, thus reducing transmission of theta activity and bursting activity, particularly as mGluR2/3 agonists have been shown to preferentially affect spike frequency ^[43,44]^.

Together, the present results demonstrate that modulating neural activity in the Papez circuit, via mGluR2 agonism, impacts early memory consolidation. Furthermore, mammillary body inhibition is sufficient to produce similar effects to those seen following systemic injections, highlighting the mammillary bodies as a vital node for consolidation. Given the mammillary bodies are affected in several neurological and psychiatric disorders^[45–52]^, this gives a greater understanding of some of the mechanisms via which disruption to the mammillary bodies can impact cognitive processing.

## Materials and Methods

### Animals

#### Rats

A total of 73 male Lister Hooded rats (Envigo, UK) were used in the study: **Experiment 1** involved 28 animals (∼320-360g at the time of injections); **Rat Experiment 3** combined 45 rats from across three separate cohorts. Rats were housed in pairs in standard large cages (**Rat Experiments 1&2**) or large Double Decker cages (**Experiment 3**, Tecniplast, UK) with enrichment (chewsticks, tunnels) in a temperature-controlled room with a 12h light/dark cycle. Prior to behavioral testing, all rats were food restricted to a minimum of 85% of their free-feeding weight. Water was available *ad libitum* throughout.

#### Mice

A total of 51 c57bl6/j mice were used: **Experiment 2** included 33 mice (15-25 weeks old at the start of the experiment, 15 female, 21 bred in-house and 12 sourced from Charles River, UK); **Experiment 3** included 18 mice (21-29 weeks of age at the start of the experiment, 3 female, all bred in-house). Mice were housed in groups of two, in a temperature-controlled room, with enrichment (chewsticks, tunnels), under a 12 h light/dark cycle and with *ad libitum* access to food and water.

All experiments were performed in accordance with the UK Animals (Scientific Procedures) Act, 1986 and associated guidelines, the EU directive 2010/63/EU, as well as the Cardiff University Biological Standards Committee; the experiments were reported according to the ARRIVE guidelines.

### Randomization and blinding

For all experiments, test parameters (e.g. arm numbers, object pairs) were counterbalanced by pseudo-randomization (Latin cross method) such that each drug treatment condition received an equivalent experience. Drug administration schedules were also pseudorandomized so that each testing day comprised vehicle and drug delivery in a roughly 50/50 ratio. For mice experiments, the different sexes were tested on separate days when possible. Experimenters were not always blind to the drug condition while running the tasks, however, there were no direct experimenter-animal interaction once the animals commenced the tasks; drug assignments were blinded at the analysis and scoring stages. For histological analyses (*c-fos* region of interest selection, cannula placement validation), as well as case exclusion based on aberrant behavioral profiles, the identity of the animals or drug assignment or test performance were obscured such that they could not inform the decisions.

### Experiment 1

Rats were subject to two behavioral testing paradigms: the open field and the novel place preference (NPP. The open field task was employed to rule out drug effects on overall activity whereas the NPP task tested the effect of the drug on memory consolidation.

For open field, 8 rats were first habituated to the apparatus over three 10 min sessions. The testing arena was a square, opaque wooden box (100 × 100 × 42cm high) lined with sawdust, located on the floor of an evenly illuminated room (280 × 295 × 260cm high). As it was a within-subject experiment, rats were tested on two separate days, approximately one week apart, receiving either eglumetad (6 mg/kg, 10.8 mM; also known as LY-354740, Tocris, UK) or saline vehicle injections, according to a counterbalanced design. Thirty mins after injections, rats were placed in the center of the open field arena for 10 min (morning session). Rats were returned to their home cages and again exposed to the open field for 10 min, 6h after the morning open field session (afternoon session). Animals were recorded with an overhead camera (GoPro Silver 7, GoPro Inc., USA) while performing the task.

The NPP task was run in a modified radial arm maze. The maze comprised a central arena (35 diameter × 25cm high) and eight radiating arms (10 × 87 × 18cm high), raised 68cm off the floor. The base of the maze was white and opaque whereas the walls of the maze were made of clear Perspex. Arm entrances were blocked off by transparent doors operated remotely by the experimenter. The tops of the arms were covered by transparent plastic sheets to prevent the rats from climbing out. A translucent foil collar was also placed around the top of the central arena to discourage rats from climbing. The maze was positioned in the center of a room (255 × 330 × 260cm high) and was surrounded by white curtains. The room was dimly lit by partially covered fluorescent batten lightbulbs affixed to the ceiling. Task performance was captured with a camera (GoPro Silver 7, GoPro Inc., USA) mounted on a boom pole above the maze.

All animals received 3-4 habituation sessions prior to testing. For habituation, rats were individually placed in the center platform for 30 s after which 2 of the 8 doors were opened and the rats were allowed to freely explore for 10 min. The two habituation arms formed a linear track and were kept the same for each rat across habituation sessions.

Each trial comprised a sample and a test phase, 6 h apart. The testing room was re-configured for each trial: curtains were decorated with ∼12 novel three-dimensional landmarks (e.g. Christmas decorations) and the center of the maze was sprayed with a novel baking aroma (e.g. vanilla, caramel, lemon, mint). This was to provide a novel context for each trial to maintain the animals’ interest. During sample runs, animals were given access to a previously unexplored set of arms 90° apart (see **Fig. 1d**). As with habituation, animals were first placed in the center arena for 30 s and then allowed to enter the two novel arms for another 10 min. Rats received intraperitoneal injections of eglumetad or saline vehicle immediately after the sample phase. They were then returned to their home cages, which were placed in a quiet room for 6 hours until test. The test phase involved animals having access to the same two arms as in the sample phase as well as access to a third novel arm at 135° from either sample arm see **Fig. 1d**). This additional arm was considered a goal arm, i.e., if animals remember the two arms from the sample phase, the goal arm should be relatively more novel and therefore more interesting. Habituation and trial arms were assigned to each rat in a pseudorandomized, counterbalanced fashion, and the experiment was run according to a within-subject design such that each rat received both injection types on separate occasions, at least 7 days apart.

*Drug occupancy*: Rats were given intraperitoneal injections of eglumetad before being returned to their home cage for a predetermined amount of time (10 min, 30 min, 1 h, 3 h, 6 h). The animals were killed by a rising concentration of CO_2_, followed by decapitation, and the brains were rapidly removed using bone scissors and forceps. A slice of brain was excised using a razor and weighed (each slice was between 100 and 150 mg for ease of homogenization), then placed inside an Eppendorf tube on dry ice. The brain samples were thawed on wet ice and ice cold 10 mM KH_2_PO_4_ *pH* buffer (*pH* = 7) was added to the tube at a 1 mg:9 ml ratio of brain sample to buffer. An electronic homogenizer was used to generate a smooth solution. The homogenized samples were then stored at -80°C until further processing.

Standard curve preparation: Seven 1:3 serial dilutions of a 10 mM dimethylsulfoxide(DMSO) stock were made, yielding an eight-point Standard Curve (SC) with the following concentrations (mM): 1, 0.33, 0.11, 0.037, 0.012, 0.0041, 0.0013, 0.00045. 1 µL of each SC dilution were dispensed into separate wells of one column of a deep welled (1 mL) 96-well polypropylene plate (Phree, Phenomenex, USA). 100 µL untreated homogenized brain was aliquoted into each well (giving final concentrations of 10, 3.3, 1.1, 0.37, 0.123, 0.041, 0.0137 and 0.0045 µM), then mixed and incubated at room temperature for 10 minutes. 300 µL of a quench solution (10 mL MeOH + 10 µL 0.1 mM carbemazapine (internal standard) DMSO stock; [final] = 0.1 µM) was added to each well and mixed. A further 600 µL MeOH was added to each well and mixed gently. The plate(s) were then centrifuged at 4000 rpm for 30 min to pellet the precipitate. The supernatant (approximately 850 µL) was then transferred into separate wells of a HybridSPE-plus 96 well filter plate (Sigma, Japan).

Sample Preparation: 100 µL of each homogenized brain sample was placed into separate wells of a deep welled (1 mL) 96-well polypropylene plate (Phree, Phenomenex, USA). 300 µL of quench solution was then added to each sample and mixed. A further 600 µL of MeOH was added to each well and mixed gently. The plates were then centrifuged at 4000 rpm for 30 min to pellet the precipitate. The supernatant (approximately 850 µL) was then transferred into separate wells of a lipid HybridSPE-plus 96 well filter plate (Sigma, Japan) taking care not to disturb the pellet.

Filtration: Experiment and standard curve samples were filtered under vacuum into a fresh 1 mL deep well, 96 well plate and placed in a vacuum concentrator (SPD120 SpeedVac Vacuum Concentrator, Thermofisher, USA). The vacuum evaporation was run for 5 hours at 50°C after which the plate was incubated overnight at room temperature. All dried samples were re-suspended in 150 µL 50% MeCN. All experimental and standard curve samples were then placed on a 96 well polypropylene plate before being sealed with a ‘zone-free’ sealing film (Alpha Laboratories, cat # ZAF-PE-50, UK) for LC/MS-MS sampling and analysis (see Supplementary Materials). All samples were analyzed in duplicate.

*mGluR2 Immunohistochemistry:* We used previously frozen/cryoprotected archival rat tissue, cut at 30 µm. Tissue was washed in PBST (0.2% Triton-X in PBS), then transferred to an EDTA buffer for 20 mins at 80°C, followed by a further 20 mins at room temperature. Endogenous peroxidase activity was blocked by incubation in 3% hydrogen peroxidase solution in methanol and unspecific binding blocked by a two-hour incubation in 3% normal horse serum (NHS) in PBST (0.2% Triton-X in PSB). Sections were incubated in mGluR2 mouse monoclonal antibody (1:500, Santa Cruz, USA, sc-271655, in PBST with 1% NHS) overnight at 4°C followed by a 2 h incubation in horse anti-mouse biotinylated secondary (1:200, Vector, 2Bscientific, UK, BA-200 in PBST with 1% NHS). The signal was amplified with an ABC kit (PK-6101 VectaStain kit, 2BScientific, UK) and developed by chromogenic peroxidase reaction (DAB Substrate Kit, 2BScientific, UK), all according to manufacturer’s instructions. Tissue sections were mounted, dehydrated, and cleared in xylene prior to coverslipping and imaging.

### Experiment 2

As with **Experiment 1**, mice were injected with either eglumetad or saline and tested on the open field and NPP paradigms in partially overlapping cohorts of 22 and 26 mice, respectively.

For open field, all mice first received three 10-min habituation sessions to the apparatus (40 cm × 40cm, white opaque square box), followed by the test session. At test, animals were injected with saline or eglumetad 30 min prior to being placed in the arena. Animals were allowed to explore the arena for 10 min and their behavior was recorded (USBFHD01M-SFV, ELP, UK). The test was carried out according to a between-subject design in a counterbalanced manner, considering injection type and the sex of the mice.

For NPP, prior to commencing the task, mice were given a minimum of three sessions to habituate them to handling and the experimental room. The task was conducted in a translucent string-and-pulley operated 8-arm radial maze (arm dimensions: length, 36 cm; width, 9 cm; height, 17 cm). The maze was positioned 100 cm above the ground and evenly illuminated by ambient light. An overhead camera (USBFHD01M-SFV, ELP, UK) was used to record behavior. The maze was surrounded by laboratory equipment (sink, fridges, recording stages), serving as visual reference cues. The task was conducted in an analogous manner to that in rats in **Experiment 1** except mice did not receive odor cues or additional spatial cues (due to the between-subject design, the mice were only run once on the task so additional cues were not needed to differentiate between test sessions) and were not habituated to the maze prior to the task [as in ^53^].

### Experiment 3

We studied the effect of intraperitoneal eglumetad injections on regional brain activity by utilizing experience-driven *c-fos* induction. Prior to the experiment, mice were habituated to the experimenter and the experimental room over three one-hour sessions involving gentle handling and placement of the home cages in the experimental room. *c-fos* expression was induced by allowing mice to individually explore a novel cross-maze (arm length: 33.5 cm, arm height: 4cm, arm width: 8 cm; made from clear Perspex, placed 64 cm from the ground) for 10 min each. The animals’ behavior was recorded using an overhead camera (USB3MP01H-BFV, ELP, UK). Six mice were injected with eglumetad and six with saline vehicle, 30 min prior to cross-maze exploration, while another six mice remained in their home cages to serve as controls. Following maze exploration, mice were returned to their home cages and euthanized 90 min later with pentobarbital overdose^[54]^. Mice underwent intracardial perfusion fixation (4% paraformaldehyde (PFA) in phosphate-buffered saline (PBS)). The brains were subsequently removed and fixed for a further 2 h in PFA followed by a 48 h incubation in 25% sucrose solution in PBS. The brains were then trimmed using a custom 3D-printed mould (courtesy of Peter Watson, Biosi, Cardiff University) and sectioned at 35 µm using a sliding stage microtome (Bright Instruments, UK). The tissue was cryopreserved at -20°C in an ethylene glycol-PBS solution until required.

For each brain region of interest, 6-8 sections were processed per mouse. Endogenous peroxidase activity was blocked by incubation in 3% hydrogen peroxidase solution in methanol and unspecific binding blocked by a two-hour incubation in 3% normal goat serum (NGS) in PBST (0.2% Triton-X in PSB). *c-fos* was detected with a rabbit monoclonal antibody (1:2,500, 9F6, Cell Signaling Technology, USA) in 1% NGS-containing PBST (incubation solution) for 72 hours at 4°C, followed by two-hour incubation in biotinylated goat anti-rabbit IgG in the incubation solution (1:200, PK-6101 VectaStain kit, 2BScientific, UK) at room temperature, amplified with an ABC kit (PK-6101 VectaStain kit, 2BScientific, UK) and developed by chromogenic peroxidase reaction with nickel enhancement (DAB Substrate Kit, 2BScientific, UK), all according to manufacturer’s instructions. Tissue sections were mounted, dehydrated, and cleared in xylene prior to coverslipping and imaging.

Whole-section images were obtained using an automated slide scanner (Olympus VS200, Olympus Life Science, Japan), at 4× magnification. The images were loaded into QuPath (v0.4.3^[55]^) and *c-fos*-expressing nuclei were automatically detected with a custom script using an image classifier (accuracy: 92%, precision: 92%, recall: 94%) trained on a subset of manually annotated sections within a predetermined set of regions of interest (ROIs). The ROIs included the medial prefrontal cortex (mPFC, combined prelimbic and infralimbic cortex), anterodorsal thalamic nucleus (AD), anteromedial thalamic nucleus (AM), anteroventral thalamic nucleus (AV), dorsal hippocampal regions: CA1, CA3 & DG, lateral (LMB) and medial mammillary body (MMB), postsubiculum (POST), granular (RSG) and dysgranular retrosplenial cortex (RSD), primary visual cortex (V1), ventral tegmental nucleus of Gudden (VTG, its *c-fos-*expressing, ventromedial part only) and dorsal subiculum (dSUB). The number of c*-fos*-positive nuclei was normalized to the area of detection (as density) and mean density values per ROI were computed for each experimental subject.

### Experiment 4

To determine the extent to which inhibition of the mammillary bodies alone, via an mGluR2/3 agonist, would impact on memory consolidation, we implanted rats with infusion cannulae and delivered the drug locally. All surgeries were performed under isoflurane anesthesia (induction 5%, maintenance 2%–2.5% isoflurane). At the time of surgery, rats weighed 256-320 g. The animals were positioned in a stereotaxic frame (David Kopf Instruments, USA) and the incisor bar adjusted to achieve a flat skull. For analgesic purposes, 0.1 ml of a 50-50 mixture of lidocaine (20mg/ml solution, Fresenius Kabi, UK) and bupivacaine (2.5mg/ml solution, Aspen Pharma, Ireland) was applied topically to the scalp and 0.05 ml Metacam (5mg/ml, Boehringer Ingelheim UK) was given subcutaneously. The scalp was incised to expose the skull and a craniotomy on the right side was made. The stereotaxic arm was set at 12° towards the midline and a single 8 mm guide cannula (Plastics One, 26 gauge, USA) targeting the medial mammillary bodies was implanted. The stereotaxic coordinates of the implant relative to bregma were anteroposterior -4.45 mm and lateral -1.95 mm. The cannula was lowered to -6.85 mm from the top of the cortex. The implant was anchored to the skull with four screws (SFE-M1.4-3-A2, Accu, UK) and held in place by bone cement (Zimmer Biomet, USA). To prevent blockages, a removable ‘dummy’ cannula (Plastics One, USA) was inserted into the guide cannula. The anterior and posterior ends of the incision were closed with sutures and the antibiotic powder Clindamycin (Pfizer, USA) was applied topically to the site. To recover, rats were given subcutaneous glucose-saline injections (5 ml) and placed in a heated chamber. This was followed by close post-operative monitoring with *ad libitum* food access for at least a week. Behavioral testing began 2-3 weeks from surgery.

Implanted rats underwent the open field and NPP tasks, as described for **Experiment 1**. For open field, rats received the drug infusion 15 min prior to testing. There was no afternoon open field session. For NPP, the drug infusion followed immediately after sample and the animals were tested 6 h later. In addition to these tasks, animals in **Experiment 4** were also tested on the object-in-place (OIP) task, which also probed the role of the mammillary bodies in memory consolidation, this time utilizing object-place relationships.

For OIP, animals were tested in a gray square wooden arena (100 × 100 × 42 cm high) with a clear acrylic lid. The floor of the arena was covered in a layer of sawdust. A cue card featuring geometric shapes was affixed to one wall of the arena to provide a polarizing cue. Additional salient visual cues, such as geometric shapes and high contrast stimuli, were attached to the walls of the testing room (280 × 295 × 260 cm high). Behavior was recorded using an overhead-mounted action camera (GoPro Silver 7, GoPro Inc, USA). The test objects were complex 3-dimensional shapes constructed from Duplo (Lego, Denmark) that varied in color and size (16 × 16 × 14 cm high - 15 × 16 × 19 cm high) and were too heavy to displace (each set comprised a red, a green, a blue and a yellow object). There were duplicates of each set of objects such that each test session used a different object set and within a session and duplicates were used for test and sample. Prior to testing, rats received five habituation sessions over 3 days. On the first day, rats were placed into the arena in home cage pairs for 10 mins in the morning and 10 mins in the afternoon. The following day, rats received two 5 min sessions individually. On the last day, individual rats received a single 5 min session. No objects were present in the arena during habituation. On test days, the task involved a sample and a test phase, separated by a 6 h delay. One object was placed in each corner of the arena, positioned 10 cm from the wall. An individual rat was transported to the room in a metal carrying box with a lid to prevent them from seeing outside the box. In the sample phase, the animal was placed in the center of the arena and allowed to explore the objects for 5 min. The rat was then individually placed in a holding cage in a quiet room for the delay period. For the 3 min test phase, identical duplicate objects were used but two of the objects swapped locations. All objects were cleaned with 70% ethanol between animals to remove odor cues and sawdust. The set of objects, the object pair that changed location, and object corner positions were counterbalanced across animals. As it was a within-subject design, all rats were tested twice, once under APDC and once under saline, this was also counterbalanced across animals. There was a minimum 7-day interval between the test sessions to maintain interest in the task.

At the conclusion of **Experiment 4**, all rats were given an overdose of pentobarbital and transcardially perfused. In some cases, we infused a fluorescent dye, acridine orange (11504047, Invitrogen, UK), to help estimate the spread of the drug infusate (Fig. 4f). Extracted brains were post-perfused in PFA for 3 hours, incubated in 25% sucrose solution for 12-24 h and cut at 40 µm on a sliding-stage microtome (Bright Instruments, UK). Mounted and dehydrated sections (1-in-4 series) were counterstained with cresyl violet to help visualize the cannula tracks. We also attempted to measure *c-fos* signal as in **Experiment 3**, however, we found cannulations produced very strong staining artifacts. While the tissue was not suitable for c-*fos* analysis, it was beneficial for identifying cannula placements (**Fig. S4b**). Brain sections were imaged using an automated slide scanner (VS200, Olympus, Japan) at 4x magnification and independently assessed by three experienced researchers blinded to the behavioral data. Cases deemed acceptable by all three researchers were included in analyses. Reasons for exclusion included: a) placement of cannula tip outside the borders of the medial mammillary nucleus, including the supramammillary nucleus, b) cannula tip bordering the third ventricle/outer edge of the mammillary bodies, c) excess mechanical damage.

### Drug administration

The drugs eglumetad (also known as LY-354740, Tocris, UK) and *2R*, *4R*-APDC (Tocris, UK, abbreviated to ‘APDC’ hereafter) were administered via intraperitoneal injection or via local brain infusion, respectively. Injections/infusions were preceded by habituation to gentle restraint, typically over 3 sessions.

For intraperitoneal eglumetad injections, mice received 10 mg/kg (at 360 µM in saline, *pH*-adjusted to within 7.0-7.4) whereas rats, 6 mg/kg (at 10.8 mM in saline, *pH*-adjusted to within 7.0-7.4) of the drug solution (a 30 g mouse would therefore receive 450 µL of solution whereas a 300 g rat, 900 µL of solution). A lower dose was used in rats based on pilot data suggesting motor impairments at higher doses. Saline was used as the vehicle control.

For infusions, rats received 0.6 µL of APDC (at 1 mM in distilled water), *pH*-adjusted to 7.0-7.4 or saline. Animals were lightly restrained, and the internal ‘dummy’ cannula was replaced by an infusion cannula (Plastics One, 33-gauge) that projected 2 mm below the tip of the guide cannula. The infusion cannula was connected via polyethylene tubing to a 10 µl Hamilton syringe mounted on an infusion pump (Harvard Apparatus, USA). A volume of 0.6µl was infused over approximately 2 min (a rate of 250 nl/min). The infusion cannula remained in place for a further 2 min to allow complete diffusion of the infusate. The ‘dummy’ cannula was re-inserted, and the animal returned to its cage.

### Data analysis

Sample sizes for consolidation experiments were informed by pilot data, which indicated that demonstrating a 75% difference in discrimination scores would require around 14 animals in a within-design experiment, given the variance in pilot cohorts. To account for cannula misplacements and case exclusions due to other factors (aberrant behavior, health issues, experimenter error), we aimed for three times as many animals in infusion experiments and twice as many animals in the between-subject mouse experiments. The primary outcome measure that confirmed sufficient sample sizes were tests of above-chance discrimination following vehicle administration.

For the open field task, the trajectories of the animals were computed in DeepLabCut^[56]^ using custom models trained in-house (for each individual experiment). The coordinates were loaded into Matlab and total distance travelled and mean distance from the center of the maze were calculated.For the NPP task, recordings of the behavioral data were manually scored without knowledge of animal assignment, to provide entry and exit times for each arm visit. Entry to an arm was defined as the full head of the animal crossing the arm threshold, followed by its entire body, while an exit was scored when the full head of the animal crossed the threshold in the opposite direction. Periods of grooming as well as excessive biting or licking of maze walls were annotated and later excluded from analysis (animals did not differ in grooming/biting behavior according to drug). The manual scores were loaded into Matlab. Using custom scripts, the number of arm entries, mean arm visit time, cumulative active exploration (sum of total arm visit times) and a cumulative discrimination index (D2) were computed. The D2 score was calculated as the difference in cumulative time spent visiting the novel arm and the mean cumulative time spent visiting familiar arms over the sum of the cumulative novel arm visit time and the mean cumulative familiar arm visit time. The index therefore ranged from -1 (exclusive preference for familiar arms), through 0 (no preference for novel or familiar arms or chance level performance) to 1 (exclusive preference for the novel arm). Positive scores signified novel place preference. D2 values were plotted as function of exploration time (1-10 min) at 1 s resolution and compared between conditions at 10 min exploration time. Both test and sample phases were analyzed, the former to ensure there were no mean group differences in animal activity levels by chance.For the OIP task, video recordings were loaded into Matlab. The position of the animal’s center of gravity was extracted and assigned to the object it was closest to. The object positions and identities were extracted based on color thresholding. Cropped video frames displaying the animal in proximity to one of the four objects were assembled into a tiff file for manual scoring in FIJI. A custom start-up macro in FIJI was written to enable button-controlled user scoring of the videos in a frame-by-frame manner. This method improves the reliability of data as the experimenter is unaware of object placement in the arena at the time of testing, as well as the experimental status of the animal, thus removing potential bias. A frame was assigned an exploration status if the nose of the animal pointed toward the object and was within 1 cm from the object’s border. Instances of biting and licking were not considered object exploration. FIJI-outputted csv files were loaded into Matlab for analysis. Total object exploration and a cumulative discrimination score (D2) were computed. The D2 score was calculated as the difference in exploration times between the displaced and non-displaced objects over the sum of total object exploration time.

For the NPP task, animals were removed from analyses if they spent less than 2 min exploring arms at either sample or test and if they idled in a single arm for longer than 1.5 min as well as in cases where running errors occurred (incorrect arms open, etc.). In total, 6 rats were excluded based on inadequate exploration levels (**Experiment 1:** two rats were excluded due to poor sample exploration and two for poor test exploration, one following eglumetad infusion and one following saline infusion; **Experiment 4**: two rats were excluded due to poor sample exploration). One saline-injected mouse was excluded from **Experiment 2** due to inadequate exploration levels at test.

For the OIP tasks, animals were removed from analyses if they failed to explore any of the objects at sample or if they spent less than 30 s exploring the objects at either sample or test and (no animals were rejected based on these criteria).

For open field /cross-maze, animals were removed from analyses if they failed to move (i.e., remained in the corner of the arena or refused to enter arms). One eglumetad-injected mouse from **Experiment 3** was removed from analyses on this basis.

### Statistical analysis

To test for the effect of drugs on behavioral performance, we utilised permutation tests appropriate to the experimental design (between-subject design: *permutationTest* ^[57]^ or within-subject design: mult_comp_perm_t1^[58]^). We used this approach as permutation testing is nonparametric and calculates exact, rather than estimated, *p*-values. Where assumptions for the equivalent parametric test are met, permutation and parametric hypothesis test probabilities are typically near identical. For experiments testing drug effects on memory, it was hypothesized eglumetad/*2R,4R*-APDC would lead to poorer performance, therefore one-sided (single-tail) tests were employed. Similarly, in open field experiments, it was hypothesized the drugs would lead to reduced motility and lower anxiety, and one-sided tests were used. Tests investigating whether animals performed above chance levels were also one-tailed since below chance performance is statistically implausible^[59]^. For any other tests (e.g., total number of arm visits on novel place preference task), we did not hypothesize a specific direction for the effect of the drug, and two-sided tests were used. To estimate the magnitude of the effects, we calculated the effect sizes in the form of Cohen’s d^[60]^, using *computeCohen_d*^[61]^, adjusted to reflect the small (<50) sample sizes used in the reported experiments.

For memory consolidation tasks, we also tested whether activity at sample explained the variance in discrimination scores observed at test. Informed by the presence of correlations between some sample metrics and discrimination at test, we built a series of linear mixed effect models (*fitlme*) incorporating total active exploration at sample (both NPP and OIP) or object discrimination (OIP only) as confounding factors. If the addition of sample data significantly improved the model (*compare* function), we also tested for a significant improvement with an interaction term (drug by confound). The most parsimonious model with the best fit was chosen in each case.

For **Experiment 3**, *c-fos* staining density was transformed to represent fold change from baseline (mean home-cage control regional density values). The following linear mixed effect model (*fitlme*) was built to test for the effect of drug injections on *c-fos* induction levels: ‘Density ∼ Drug × Region + (1|Animal)’. Simple effects were investigated in an analogous manner, for each individual ROI (‘Density ∼ Drug + (1|Animal)’) and the computed regional *p-*values were corrected for with the Benjamini-Hochberg method *fdr_bh* add-on^[62]^.

Manual scoring of behavioral data was performed by several experienced researchers who were unaware of the animals’ experimental status at the time of scoring. To ascertain the robustness of the scores, a sub-sample of the videos was re-scored by another researcher. For the NPP task, we found an interclass correlation coefficient (icc) of 0.90 (CI: 0.78, 0.95); *icc21* function^[63]^.

All statistical analyses were performed with custom scripts in Matlab (2022b, MathWorks, USA), available upon request. All plots were made with *gramm*^[64]^ in Matlab and modified in PowerPoint (Microsoft, USA).

## Acknowledgements

This work was funded by a Wellcome Trust Senior Research Fellowship awarded to SDV (WT212273/Z/18). SH is funded by a BBSRC PhD studentship. Thanks to Marcus Hanley from the Medicines Discovery Institute, Cardiff University, for carrying out brain occupancy analyses and to Peter Watson from BIOSI, Cardiff University for help with 3D printed brain matrices.

## Author contributions

MMM and JP performed the surgeries. MMM, JP, TH, SH, EA carried out data collection and behavioral scoring. MMM wrote the code for data analysis, prepared the figures and carried out statistical analyses. MMM, JCP and SDV wrote the main manuscript text. All authors reviewed the manuscript.

## Data availability statement

The datasets generated during the current study as well as code used in analysis are available from the corresponding author on request.

## Supplementary Materials

### Supplementary Figures

**Fig. S1.**
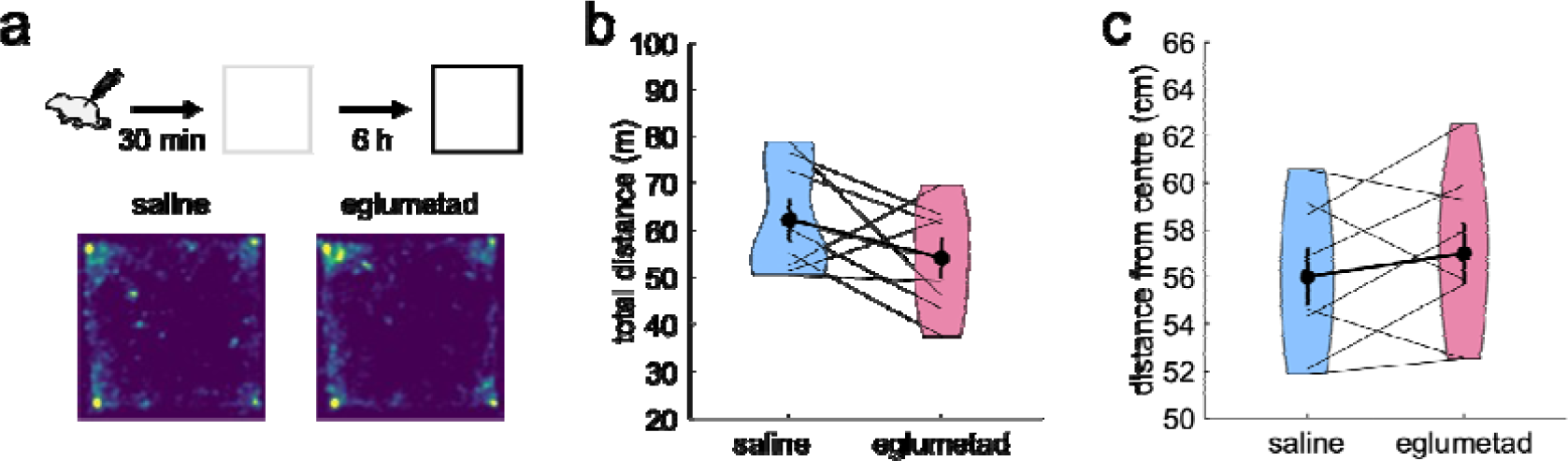
The effect of eglumetad injections on open field after a 6 h delay. **a** – Schematic representation of the task and heatmaps showing mean arena occupancy upon a second exposure to the arena 6.5 h from injection (n = 8, within-subject design). **b&c** – Violin plots of the total distance travelled and mean distance from the center of the maze, respectively (as in **Fig. 1 b&c**).

**Fig. S2.**
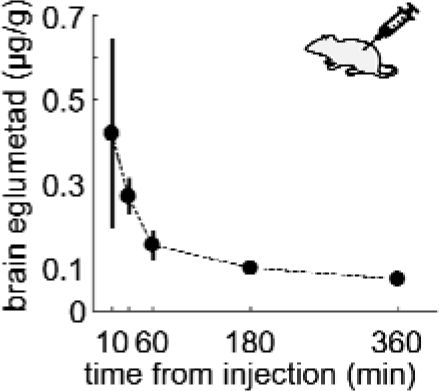
Concentration of eglumetad in rat brain at increasing intervals from intraperitoneal injection (10 min, 30 min, 1 h, 3 h, 6 h). The dots represent means and the vertical bars, the standard error of the mean (n = 20 with one rat serving as blank control).

**Fig. S3.**
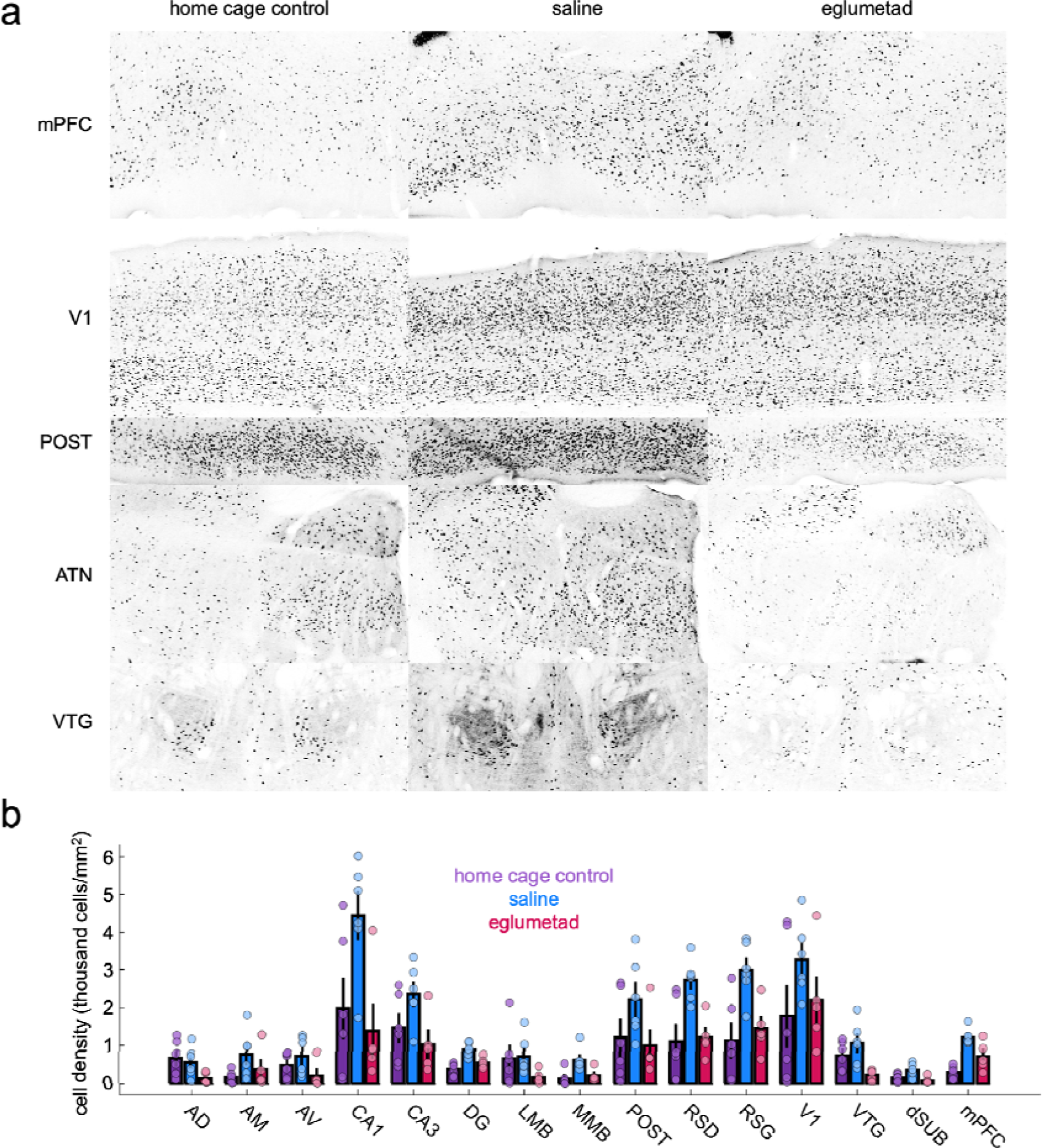
Raw c-fos expression levels in home. **a** – Representative images of regional c-fos expression levels in home cage controls, saline-injected and eglumetad-injected mice. **b** – Quantification of regional cell densities in home cage controls, saline-injected and eglumetad-injected mice. Abbreviations as in **Fig. 3**.

**Fig. S4.**
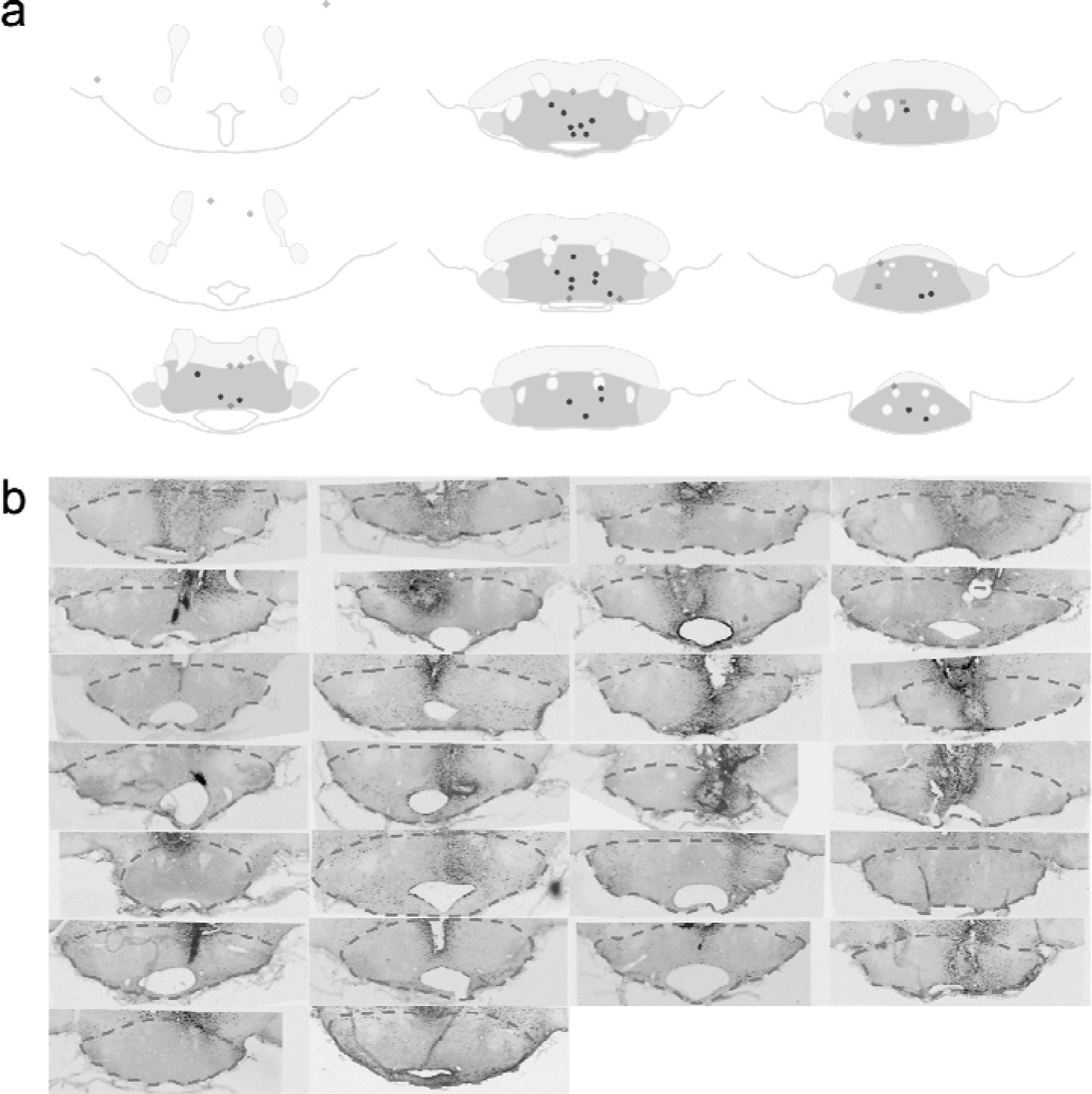
Histological validation of local drug infusion. **a** – Estimated positions of cannula tips across 9 coronal levels. **b** – Images showing cannula tip location. The dark hyperintensities are caused by heightened DAB staining in scarred tissue around implants.

**Fig. S5.**
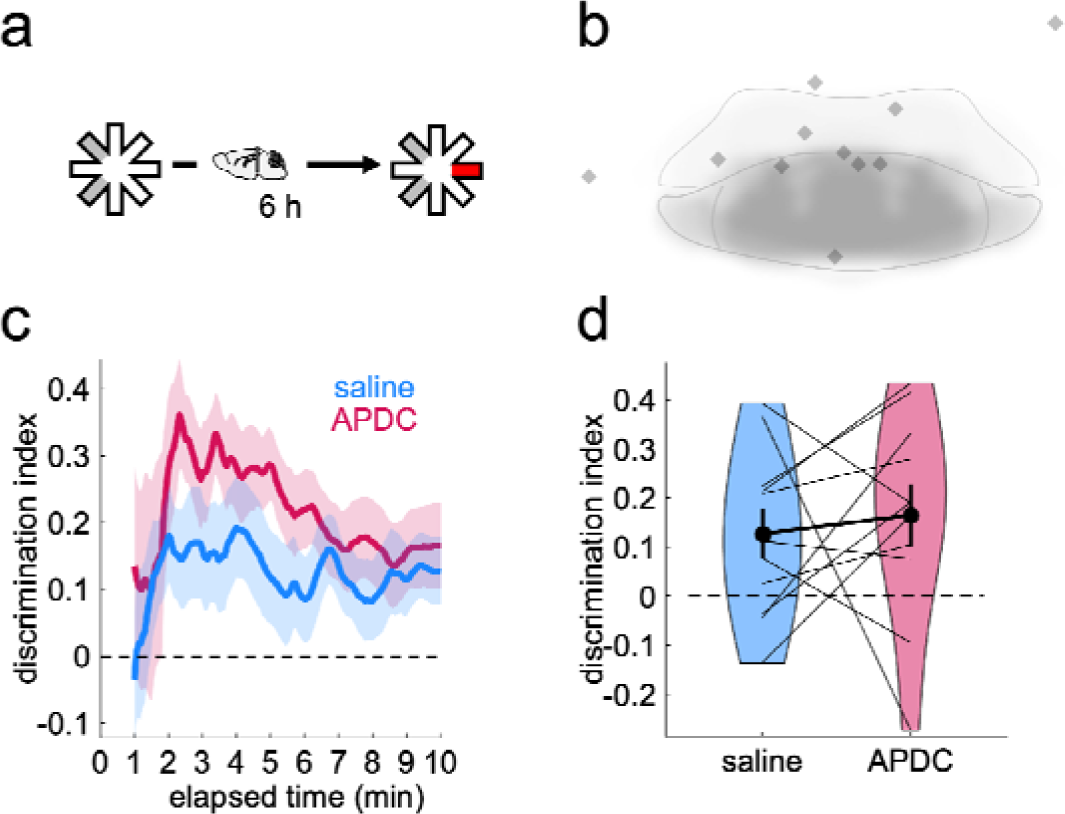
The absence of drug effect in miscannulated animals. **a** – Schematic representation of the novel place preference task. **b** – Estimated positions of misplaced cannulae. Note, due to fiber encapsulation of the mammillary bodies, infusions made at the dorsal boundaries do not spread into the mammillary bodies. **c** – Timeline of cumulative discrimination for miscannulated animals (n = 11; excludes two cases removed from analyses due to poor sample exploration activity, see Methods). **d** – Violin plots of cumulative discrimination scores at 10 min of test exploration.

## Supplementary Methods for Brain Occupancy Analysis

Liquid chromatography–mass spectrometry analysis: Analyses were carried out using the Acquity UPLC system (Waters Corporation, USA). 20 µL of each sample (tested in duplicate) were subjected to the following chromatography: Solvent A - Milli-Q water + 0.2% Acetonitrile + 0.1 % formic acid (F.A.); Solvent B – Acetonitrile + 0.1 % F.A.; Flow rate – 0.5 mL/min; Column (Waters HSS T3, 2.1 × 50 mm, 1.8 µm particle); Guard Column - ACQUITY UPLC HSS T3 VanGuard Pre-column, 100Å, 1.8 µm, 2.1 mm × 5 mm, (Waters, cat # 186003976).

Gradient (5 min):

| Time (min) | Solvent A % | Solvent B % | Curve |
| --- | --- | --- | --- |
| 0 | 100 | 0 | 1 |
| 0.5 | 100 | 0 | 11 |
| 2.5 | 0 | 100 | 6 |
| 5 | 100 | 0 | 1 |

### Mass spectrometry analysis

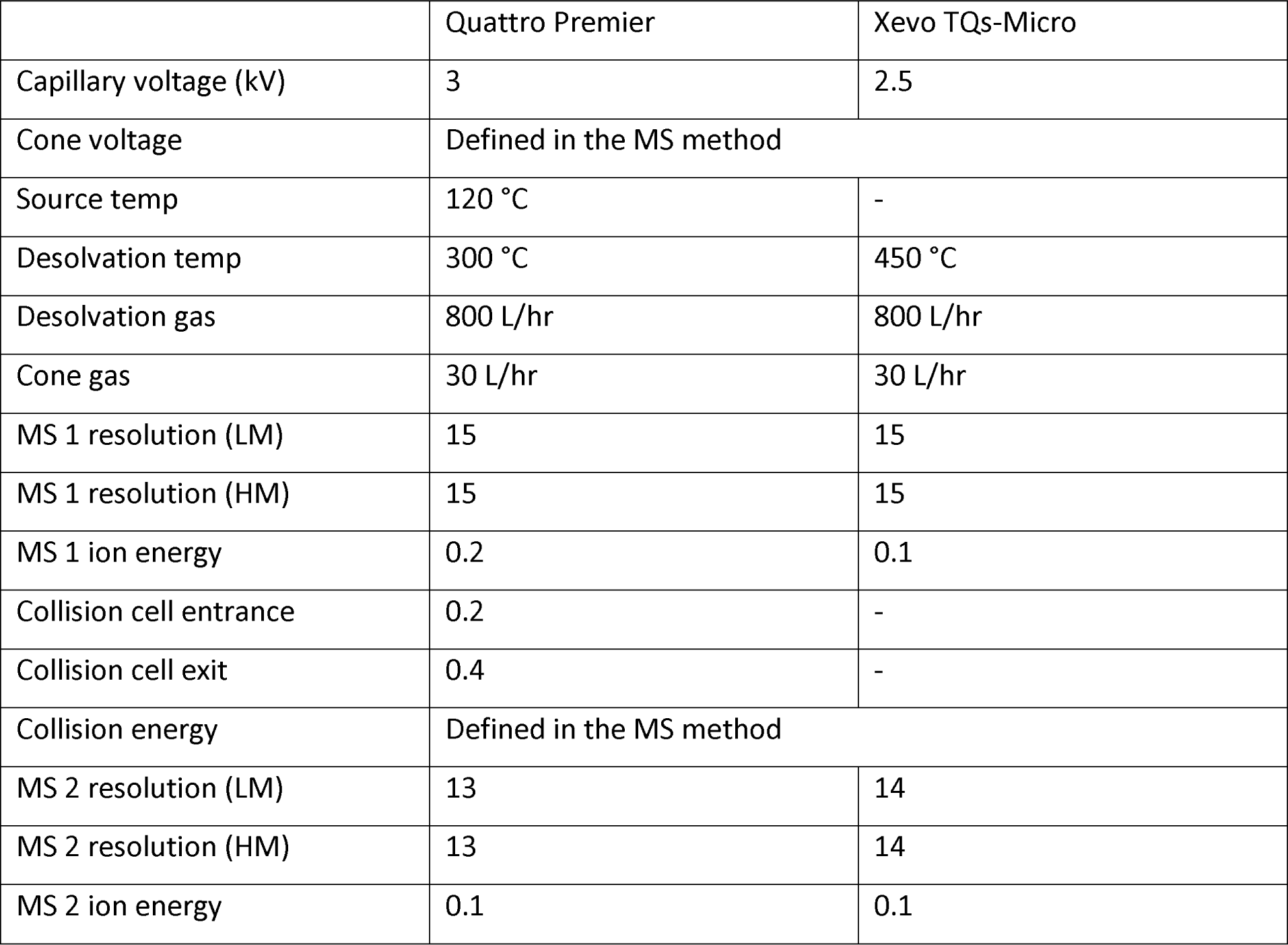

## References

1 Danet, L. et al. Thalamic amnesia after infarct. Neurology 85, 2107, doi:10.1212/WNL.0000000000002226 (2015).

2 McNaughton, N. & Vann, S. D. Construction of complex memories via parallel distributed cortical-subcortical iterative integration. Trends Neurosci. 45, 550–562, doi:10.1016/j.tins.2022.04.006 (2022).

3 Nelson, A. J. & Vann, S. D. Mammilliothalamic tract lesions disrupt tests of visuo-spatial memory. Behav. Neurosci. 128, 494–503, doi:10.1037/bne0000001 (2014).

4 Park, K. C. et al. Amnesic syndrome in a mammillothalamic tract infarction. J Korean Med Sci 22, 1094–1097, doi:10.3346/jkms.2007.22.6.1094 (2007).

5 Vann, S. D. & Aggleton, J. P. Evidence of a spatial encoding deficit in rats with lesions of the mammillary bodies or mammillothalamic tract. J. Neurosci. 23, 3506–3514, doi:23/8/3506 [pii] (2003).

6 Wetzel, C. D. & Squire, L. R. Encoding in anterograde amnesia. Neuropsychologia 18, 177–184 (1980).

7 Cermak, L. S., Uhly, B. & Reale, L. Encoding specificity in the alcoholic Korsakoff patient. Brain Lang. 11, 119–127 (1980).

8 Parkin, A. J. in Neuropsychology of Memory (eds L. R. Squire & N. Butters) (Guildford Press, 1992).

9 Squire, L. R. Two forms of human amnesia: an analysis of forgetting. J. Neurosci. 1, 635–640 (1981).

10 Atherton, L. A., Dupret, D. & Mellor, J. R. Memory trace replay: the shaping of memory consolidation by neuromodulation. Trends Neurosci. 38, 560–570, doi:10.1016/j.tins.2015.07.004 (2015).

11 Samanta, A., Alonso, A. & Genzel, L. Memory reactivations and consolidation: considering neuromodulators across wake and sleep. Current Opinion in Physiology 15, 120–127, 10.1016/j.cophys.2020.01.003 (2020).

12 Klinzing, J. G., Niethard, N. & Born, J. Mechanisms of systems memory consolidation during sleep. Nat. Neurosci. 22, 1598–1610, doi:10.1038/s41593-019-0467-3 (2019).

13 Dillingham, C. M. et al. Mammillothalamic disconnection alters hippocampocortical oscillatory activity and microstructure: Implications for diencephalic amnesia. J. Neurosci. 39, 6696–6713, doi:10.1523/jneurosci.0827-19.2019 (2019).

14 Vann, S. D. Re-evaluating the role of the mammillary bodies in memory. Neuropsychologia 48, 2316–2327, doi:S0028-3932(09)00422-9 [pii]10.1016/j.neuropsychologia.2009.10.019 (2010).

15. Dillingham, C. M., Wilson, J. J. & Vann, S. D. Electrophysiological properties of the medial mammillary bodies across the sleep-wake cycle. bioRxiv, 2023.2010.2030.563083, doi:10.1101/2023.10.30.563083 (2023).

16 Tambini, A. & Davachi, L. Awake Reactivation of Prior Experiences Consolidates Memories and Biases Cognition. Trends Cogn Sci 23, 876–890, doi:10.1016/j.tics.2019.07.008 (2019).

17 Żakowski, W. & Zawistowski, P. Neurochemistry of the mammillary body. Brain Struct Funct 228, 1379–1398, doi:10.1007/s00429-023-02673-4 (2023).

18 Hou, Y. et al. Topographical organization of mammillary neurogenesis and efferent projections in the mouse brain. Cell Reports 34, 108712, 10.1016/j.celrep.2021.108712 (2021).

19 Ohishi, H., Neki, A. & Mizuno, N. Distribution of a metabotropic glutamate receptor, mGluR2, in the central nervous system of the rat and mouse: an immunohistochemical study with a monoclonal antibody. Neurosci. Res. 30, 65–82, 10.1016/S0168-0102(97)00120-X (1998).

20 Lyon, L. et al. Fractionation of Spatial Memory in GRM2/3 (mGlu2/mGlu3) Double Knockout Mice Reveals a Role for Group II Metabotropic Glutamate Receptors at the Interface Between Arousal and Cognition. Neuropsychopharmacology 36, 2616–2628, doi:10.1038/npp.2011.145 (2011).

21 Schlumberger, C. et al. Effects of a metabotropic glutamate receptor group II agonist LY354740 in animal models of positive schizophrenia symptoms and cognition. Behavioural Pharmacology 20 (2009).

22 Higgins, G. A. et al. Pharmacological manipulation of mGlu2 receptors influences cognitive performance in the rodent. Neuropharmacology 46, 907–917, 10.1016/j.neuropharm.2004.01.018 (2004).

23 Griebel, G. et al. The mGluR2 positive allosteric modulator, SAR218645, improves memory and attention deficits in translational models of cognitive symptoms associated with schizophrenia. Scientific Reports 6, 35320, doi:10.1038/srep35320 (2016).

24 Wolf, D. H. et al. Effect of mGluR2 positive allosteric modulation on frontostriatal working memory activation in schizophrenia. Mol. Psychiatry 27, 1226–1232, doi:10.1038/s41380-021-01320-w (2022).

25 Riedel, G., Platt, B. & Micheau, J. Glutamate receptor function in learning and memory. Behav. Brain Res. 140, 1–47, 10.1016/S0166-4328(02)00272-3 (2003).

26 Crupi, R., Impellizzeri, D. & Cuzzocrea, S. Role of Metabotropic Glutamate Receptors in Neurological Disorders. Front Mol Neurosci 12, doi:10.3389/fnmol.2019.00020 (2019).

27 Nadia, R., Urs, G. & Jeanne, S. Activation of Group II Metabotropic Glutamate Receptors Promotes LTP Induction at Schaffer Collateral-CA1 Pyramidal Cell Synapses by Priming NMDA Receptors. The Journal of Neuroscience 36, 11521, doi:10.1523/JNEUROSCI.1519-16.2016 (2016).

28 Ahnaou, A., Ver Donck, L. & Drinkenburg, W. H. I. M. Blockade of the metabotropic glutamate (mGluR2) modulates arousal through vigilance states transitions: Evidence from sleep–wake EEG in rodents. Behav. Brain Res. 270, 56–67, 10.1016/j.bbr.2014.05.003 (2014).

29 Lee, C. C. Inhibition of mammillary body neurons by direct activation of Group II metabotropic glutamate receptors. Neurotransmitter (Houst) 3 (2016).

30 Lam, A. G., Monn, J. A., Schoepp, D. D., Lodge, D. & McCulloch, J. Group II selective metabotropic glutamate receptor agonists and local cerebral glucose use in the rat. J Cereb Blood Flow Metab 19, 1083–1091, doi:10.1097/00004647-199910000-00004 (1999).

31 David, R. H., Joseph, P. T., James, A. M., Darryle, D. S. & Mary Jeanne, K. Anxiolytic and Side-Effect Profile of LY354740: A Potent, Highly Selective, Orally Active Agonist for Group II Metabotropic Glutamate Receptors. J. Pharmacol. Exp. Ther. 284, 651 (1998).

32 Parker, A. & Gaffan, D. Mamillary Body Lesions in Monkeys Impair Object-in-Place Memory: Functional Unity of the Fornix-Mamillary System. J. Cognit. Neurosci. 9, 512–521, doi:doi:10.1162/jocn.1997.9.4.512 (1997).

33 Gromova, E. A., Plakkhinas, L. A. & Nesterova, I. V. [The role of the mammillary complex in consolidating memory traces]. Zhurnal vysshei nervnoi deiatelnosti imeni I P Pavlova 25, 372–378 (1975).

34 Sif, J. et al. Time-dependent sequential increases in [14C]2-deoxyglucose uptake in subcortical and cortical structures during memory consolidation of an operant training in mice. Behav Neural Biol 56, 43–61, doi:10.1016/0163-1047(91)90279-y (1991).

35 Bontempi, B., Jaffard, R. & Destrade, C. Differential temporal evolution of post-training changes in regional brain glucose metabolism induced by repeated spatial discrimination training in mice: visualization of the memory consolidation process? Eur. J. Neurosci. 8, 2348–2360, doi:10.1111/j.1460-9568.1996.tb01198.x (1996).

36 Mendez-Lopez, M., Mendez, M., Sampedro-Piquero, P. & Arias, J. L. Spatial learning-related changes in metabolic activity of limbic structures at different posttask delays. J. Neurosci. Res. 91, 151–159, doi:10.1002/jnr.23134 (2013).

37 Melo, M. B. d., Favaro, V. M. & Oliveira, M. G. M. The dorsal subiculum is required for contextual fear conditioning consolidation in rats. Behav. Brain Res. 390, 112661, 10.1016/j.bbr.2020.112661 (2020).

38 Safari, V. et al. Individual Subnuclei of the Rat Anterior Thalamic Nuclei Differently affect Spatial Memory and Passive Avoidance Tasks. Neuroscience 444, 19–32, 10.1016/j.neuroscience.2020.07.046 (2020).

39 Skelin, I., Kilianski, S. & McNaughton, B. L. Hippocampal coupling with cortical and subcortical structures in the context of memory consolidation. Neurobiol. Learn. Mem. 160, 21–31, 10.1016/j.nlm.2018.04.004 (2019).

40 Li, M. L., Hu, X. Q., Li, F. & Gao, W. J. Perspectives on the mGluR2/3 agonists as a therapeutic target for schizophrenia: Still promising or a dead end? Prog Neuropsychopharmacol Biol Psychiatry 60, 66–76, doi:10.1016/j.pnpbp.2015.02.012 (2015).

41 Petralia, R. S., Wang, Y. X., Niedzielski, A. S. & Wenthold, R. J. The metabotropic glutamate receptors, MGLUR2 and MGLUR3, show unique postsynaptic, presynaptic and glial localizations. Neuroscience 71, 949–976, 10.1016/0306-4522(95)00533-1 (1996).

42 Jin, L. E. et al. mGluR2/3 mechanisms in primate dorsolateral prefrontal cortex: evidence for both presynaptic and postsynaptic actions. Mol. Psychiatry 22, 1615–1625, doi:10.1038/mp.2016.129 (2017).

43 Keele, N. B., Neugebauer, V. & Shinnick-Gallagher, P. Differential Effects of Metabotropic Glutamate Receptor Antagonists on Bursting Activity in the Amygdala. J. Neurophysiol. 81, 2056–2065, doi:10.1152/jn.1999.81.5.2056 (1999).

44 Kintscher, M., Breustedt, J., Miceli, S., Schmitz, D. & Wozny, C. Group II metabotropic glutamate receptors depress synaptic transmission onto subicular burst firing neurons. PLoS One 7, e45039, doi:10.1371/journal.pone.0045039 (2012).

45 Molavi, M., Vann, S. D., de Vries, L. S., Groenendaal, F. & Lequin, M. Signal Change in the Mammillary Bodies after Perinatal Asphyxia. AJNR. American journal of neuroradiology 40, 1829–1834, doi:10.3174/ajnr.A6232 (2019).

46 Milczarek, M. M., Gilani, S. I. A., Lequin, M. H. & Vann, S. D. Reduced mammillary body volume in individuals with a schizophrenia diagnosis: an analysis of the COBRE data set. Schizophrenia (Heidelb) 9, 48, doi:10.1038/s41537-023-00376-7 (2023).

47 Meys, K. M. E., de Vries, E., Groenendaal, F., Vann, S. & Lequin, M. H. The mammillary bodies: a review of causes of injury in infants and children. AJNR. American journal of neuroradiology, doi:10.3174/ajnr.A7463 (2022).

48 Lequin, M. H. et al. Mammillary body injury in neonatal encephalopathy: a multicentre, retrospective study. Pediatr Res, doi:10.1038/s41390-021-01436-3 (2021).

49 Connaughton, M. et al. The Limbic System in Children and Adolescents with Attention-Deficit/Hyperactivity Disorder: A Longitudinal Structural MRI Analysis. Biological Psychiatry Global Open Science, 10.1016/j.bpsgos.2023.10.005 (2023).

50 Denby, C. E. et al. The frequency and extent of mammillary body atrophy associated with surgical removal of a colloid cyst. AJNR. American journal of neuroradiology 30, 736–743, doi:ajnr.A1424 [pii] 10.3174/ajnr.A1424 (2009).

51 Bernstein, H. G. et al. A postmortem assessment of mammillary body volume, neuronal number and densities, and fornix volume in subjects with mood disorders. Eur Arch Psychiatry Clin Neurosci 262, 637–646, doi:10.1007/s00406-012-0300-4 (2012).

52 Küçükerden, M. et al. Compromised mammillary body connectivity and psychotic symptoms in mice with di- and mesencephalic ablation of ST8SIA2. Translational Psychiatry 12, 51, doi:10.1038/s41398-022-01816-1 (2022).

53 Nicole, O. et al. Soluble amyloid beta oligomers block the learning-induced increase in hippocampal sharp wave-ripple rate and impair spatial memory formation. Scientific Reports 6, 22728, doi:10.1038/srep22728 (2016).

54 Morgan, J. I., Cohen, D. R., Hempstead, J. L. & Curran, T. Mapping Patterns of c-*fos* Expression in the Central Nervous System After Seizure. Science 237, 192–197, doi:doi:10.1126/science.3037702 (1987).

55 Bankhead, P. et al. QuPath: Open source software for digital pathology image analysis. Scientific Reports 7, 16878, doi:10.1038/s41598-017-17204-5 (2017).

56 Mathis, A. et al. DeepLabCut: markerless pose estimation of user-defined body parts with deep learning. Nat. Neurosci. 21, 1281–1289, doi:10.1038/s41593-018-0209-y (2018).

57 Krol, L. R. Permutation Test (https://github.com/lrkrol/permutationTest), GitHub. Retrieved November 13, 2023. (2023).

58 Groppe, D. mult_comp_perm_t1(data,n_perm,tail,alpha_level,mu,reports,seed_state) (https://www.mathworks.com/matlabcentral/fileexchange/29782-mult_comp_perm_t1-data-n_perm-tail-alpha_level-mu-reports-seed_state), MATLAB Central File Exchange. Retrieved November 13, 2023. (2023).

59 Akkerman, S., Prickaerts, J., Steinbusch, H. W. M. & Blokland, A. Object recognition testing: Statistical considerations. Behav. Brain Res. 232, 317–322, 10.1016/j.bbr.2012.03.024 (2012).

60 Cohen, J. Statistical Power Analysis for the Behavioral Sciences. 2nd Edition edn, (Lawrence Erlbaum Associates, Publishers, 1988).

61 Bettinardi, R. G. computeCohen_d(x1, x2, varargin) (https://www.mathworks.com/matlabcentral/fileexchange/62957-computecohen_d-x1-x2-varargin), MATLAB Central File Exchange. Retrieved November 13, 2023. (2023).

62 Groppe, D. fdr_bh (https://www.mathworks.com/matlabcentral/fileexchange/27418-fdr_bh), MATLAB Central File Exchange. Retrieved November 13, 2023. (2023).

63 Zoeller, T. Intraclass correlation coefficient with confidence intervals (https://www.mathworks.com/matlabcentral/fileexchange/26885-intraclass-correlation-coefficient-with-confidence-intervals), MATLAB Central File Exchange. Retrieved November 14, 2023. (2023).

64 Morel, P. Gramm: grammar of graphics plotting in Matlab. Journal of Open Source Software 3, 10.21105/joss.00568 (2018).

